# Nanoscale CaV channel reorganization links α-synuclein pathology to calcium-dependent transcriptional dysregulation

**DOI:** 10.64898/2026.05.01.719272

**Authors:** Zoltán J. Kovács, Rose Ellen Dixon, Eamonn James Dickson

**Affiliations:** Department of Physiology and Membrane Biology, University of California, Davis, California, 95616

**Keywords:** Alpha-Synuclein (α-Synuclein), alpha-Synuclein pre-formed fibrils (α-Synuclein PFF), Cyclin Dependent Kinase 5 (CDK5), Parkinson’s Disease (PD), Voltage-gated calcium channel 1.2 (Ca_V_1.2), Voltage-gated calcium channel 2.1 (Ca_V_2.1), Voltage-gated potassium channel 2.1 (K_V_2.1)

## Abstract

Disrupted calcium (Ca^2+^) homeostasis is a hallmark of neurodegenerative diseases, yet the mechanisms driving excessive Ca^2+^ entry at the neuronal plasma membrane remain poorly understood. Here, we show that α-synuclein pre-formed fibrils trigger a reorganization of voltage-gated calcium (Ca_V_) channels in cultured mouse cortical neurons, increasing their clustering at the soma and dendrites. We find that α-synuclein fibrils promote cyclin-dependent kinase 5–mediated phosphorylation of K_V_2.1 at serine 603, enhancing its scaffolding capacity for Ca_V_ channels. Disrupting Ca_V_–K_V_2.1 coupling with a competitive peptide or pharmacological CDK5 inhibition restores channel proximity to control levels. Functionally, the enhanced Ca_V_ clustering amplifies depolarization-evoked Ca^2+^ influx and drives aberrant excitation–transcription coupling, as reflected by elevated expression of the immediate early gene c-Fos. All effects were rescued by disrupting Ca_V_–K_V_2.1 interactions or inhibiting Ca_V_ channel activity. These findings identify nanoscale Ca_V_ channel remodeling as a mechanistic link between α-synuclein pathology and Ca^2+^-dependent transcriptional dysregulation, positioning Parkinson’s disease as a nanostructural channelopathy.

## Introduction

α-synuclein is a small, abundantly expressed neuronal protein that coordinates synaptic vesicle trafficking, neurotransmitter release, organelle homeostasis, and activity-dependent signaling through dynamic interactions with diverse protein partners and membranes ^1–9^. In synucleinopathies such as Parkinson’s disease (PD), α-synuclein misfolds and assembles into toxic oligomeric and fibrillar species that propagate between neurons and accumulate as Lewy bodies, the pathological hallmark of PD ^6,10,11^. This incurable, progressive neurodegenerative disorder is characterized by motor impairments, including tremor, bradykinesia, and rigidity, together with non-motor symptoms such as cognitive decline, with 85–90% of cases being idiopathic in origin ^12,13^. α-synuclein fibrils exert multiple neurotoxic effects through disruption of membrane integrity, impaired vesicle trafficking, and mitochondrial dysfunction ^6,14,15^, yet how they disrupt calcium (Ca^2+^) homeostasis, a key regulator of neuronal excitability, synaptic transmission, intracellular signaling, and cell survival, remains incompletely understood^16^.

Current models emphasize that α-synuclein disrupts ER Ca^2+^ handling, promoting excessive IP_3_ receptor–mediated release and pathological Ca^2+^ transfer to mitochondria at ER–mitochondria contact sites ^9,17,18^. However, the ER is a finite store, and sustained release requires replenishment from the extracellular space. Which plasma membrane Ca^2+^ entry pathway(s) is remodeled by α-synuclein pathology remains an open question.

Voltage-gated calcium channels (Ca_V_) are strong candidates for mediating this entry. Ca_V_ channels provide the principal route for depolarization-evoked Ca^2+^ influx across the neuronal plasma membrane and play essential roles in neurotransmission, synaptic plasticity, and the activation of Ca^2+^-dependent signaling cascades^19^. Importantly, Ca_V_ channels are not uniformly distributed but can assemble into nanoscale clusters at the plasma membrane, an organization that increases local channel density, promotes cooperative gating, and amplifies channel activity ^20–23^. Conversely, excessive Ca_V_ hyperactivity have been implicated in neuronal dysfunction across several neurological disorders ^24–28^, while pharmacological Ca_V_ inhibition confers neuroprotection in models of PD ^29–31^.

In the neuronal soma, prominent Ca_V_ clusters are concentrated at ER–plasma membrane (ER–PM) junctions organized by clustered K_V_2.1 potassium channels that tether to ER-resident VAP proteins^32^. K_V_2.1 clustering is tightly regulated by its phosphorylation state: kinase-mediated phosphorylation promotes clustering in a non-conducting scaffold configuration^33–35^, whereas phosphatase-driven dephosphorylation disperses clusters and restores channel conductance ^36,37^. At these junctions, clustered K_V_2.1 channels recruit and stabilize Ca_V_ channels, while ER-resident ryanodine receptors (RyRs) contribute local Ca^2+^ release, creating Ca^2+^ signaling hotspots ^28,38,39^. Ca_V_ channels at these sites are also established mediators of excitation-transcription coupling, where activity-dependent Ca^2+^ influx activates intracellular signaling pathways that drive gene expression^38^. Given that transcriptomic studies have revealed widespread gene expression alterations in PD ^40^, Ca_V_ channels at K_V_2.1-organized junctions emerge as compelling candidates linking α-synuclein-driven Ca^2+^ imbalance to transcriptional dysregulation.

Here, we show that human α-synuclein pre-formed fibrils induce a nanoscale reorganization of Ca_V_ channels at the neuronal plasma membrane through a CDK5-dependent increase in K_V_2.1 phosphorylation that enhances K_V_2.1 scaffolding and promotes Ca_V_ retention and clustering. This enhanced Ca_V_ clustering amplifies depolarization-evoked Ca^2+^ influx and drives aberrant Ca^2+^-dependent gene expression. Importantly, disrupting Ca_V_–K_V_2.1 coupling normalizes Ca^2+^ entry, while pharmacological Ca_V_ inhibition abrogates dysregulated gene expression, identifying Ca_V_ channel remodeling as both a mechanistic link between α-synuclein pathology and transcriptional dysregulation and a potential therapeutic target

## Results

### α-synuclein pre-formed fibrils increase Ca_V_ clustering at the neuronal plasma membrane

To investigate the effects of α-synuclein pathology on Ca_V_ channel organization, we treated cultured mouse cortical neurons with human α-synuclein Type I pre-formed fibrils (hereafter referred to as PFFs) (**Fig. S1A**). Neurons internalize PFFs via endocytic pathways, where the fibrils recruit endogenous α-synuclein into aberrant, insoluble aggregates ^6,41^. Because this model is independent of α-synuclein mutations, it provides a platform to study the more prevalent idiopathic form of PD ^6,41–45^. While dopaminergic neurons of the substantia nigra are the most vulnerable population in early PD, α-synuclein pathology progressively spreads to cortical regions as the disease advances ^14,41,43,46^, a process associated with severe cognitive decline and dementia. Cortical neurons express α-synuclein and are susceptible to Lewy body formation^47–50^, making them a relevant model for studying the mechanisms of α-synuclein-driven neuronal dysfunction in later disease stages. We first confirmed robust neuronal uptake of ATTO594-labeled PFFs and found that a single fibril treatment is sufficient to achieve efficient internalization (**Fig. S1B**). Therefore, all subsequent experiments used a single treatment with unlabeled PFFs (**Fig. S1A**).

The association of α-synuclein with disrupted Ca^2+^ homeostasis^51^, together with evidence that Ca_V_ clusters undergo disease-related remodeling^52–54^, prompted us to examine whether PFFs alter Ca_V_ organization. Neurons were treated with unlabeled PFFs (see Methods), fixed, and immunolabeled for Ca_V_1.2, a major L-type calcium channel widely expressed in the cortex^25^. To enable cell-type-specific analysis, neurons were co-stained for the inhibitory marker glutamic acid decarboxylase GAD-67^55^, allowing us to distinguish Ca_V_1.2 clustering in excitatory and inhibitory neurons (**Fig. S1C**). Quantitative analysis of super-resolution (∼120 nm resolution^56^) confocal images acquired from somatic PM regions (**Fig. 1A**) revealed that PFF treatment significantly increased Ca_V_1.2 cluster size, density, and mean gray value (MGV) (**Fig. 1B**). In dendrites, cluster size and MGV were also significantly increased in both excitatory and inhibitory neurons (**Fig. 1B).** To independently validate these findings at higher resolution, we generated localization maps using super-resolution single-molecule total internal reflection fluorescence (TIRF) microscopy (resolution ∼20 nm^57,58^). These maps confirmed that PFF treatment increased both the size and density of PM Ca_V_1.2 clusters in the soma (**Fig. 1C**). Nearest neighbor distance analysis further revealed that PFF exposure reduced the inter-cluster spacing between adjacent Ca_V_1.2 clusters (**Fig. 1C**). Because the observed increase in Ca_V_1.2 clustering could arise either from elevated protein expression or from redistribution of channels from intracellular endomembrane reserves to the plasma membrane, we assessed total Ca_V_1.2 expression levels by western blot (WB). No significant change was detected following PFF treatment, indicating that PFF exposure does not affect overall Ca_V_1.2 expression but instead modulates its distribution within the plasma membrane (**Fig. S1D**).

**Fig. 1.**
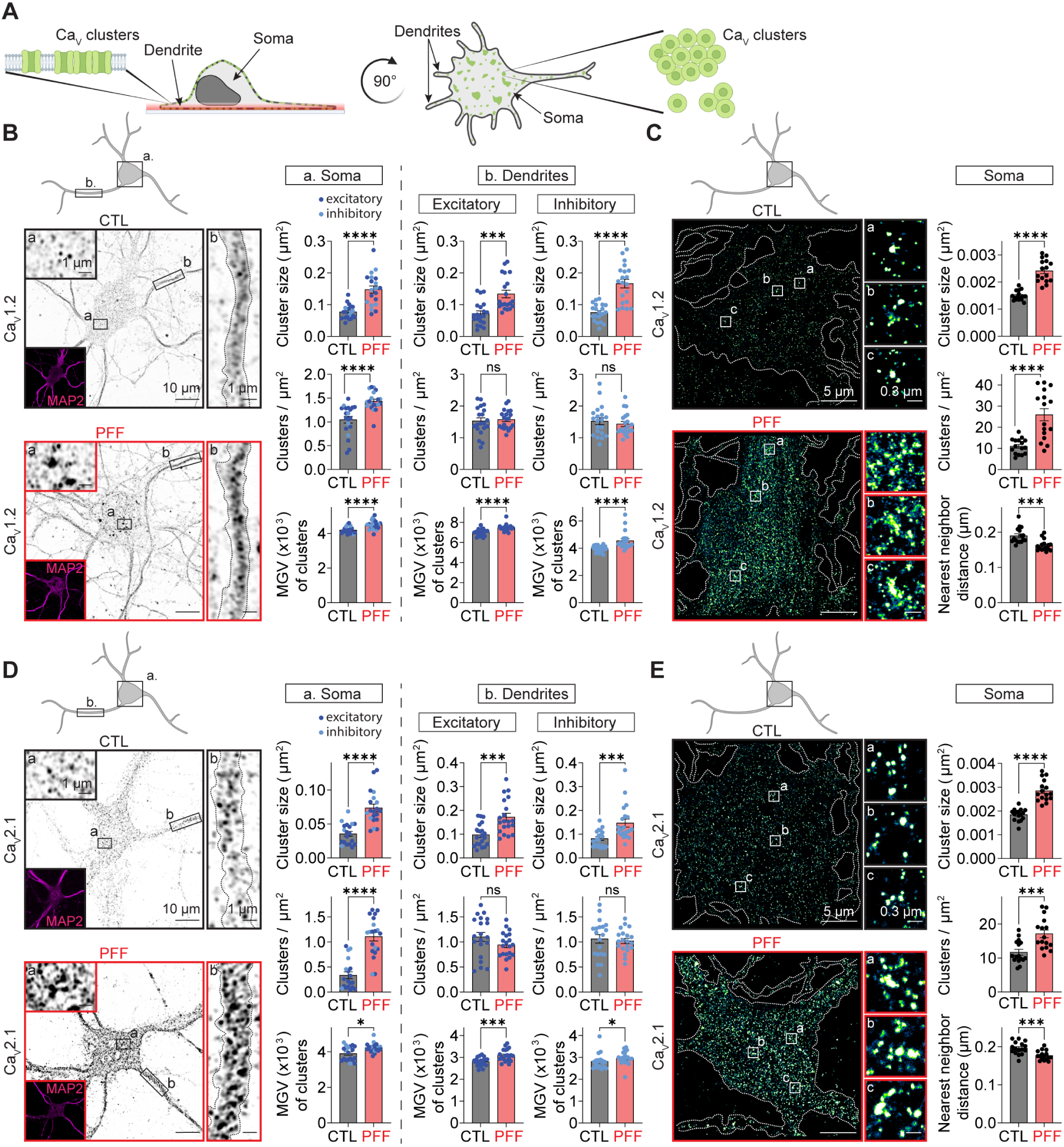
PFF treatment increases Ca_V_ channel clustering at the neuronal plasma membrane. **A** Schematic illustrating imaging at the plasma membrane (PM). **B** Left: representative single-plane Airyscan confocal images of the PM showing Ca_V_1.2 immunolabeling in control (CTL, black) and PFF-treated (red) neurons. Inset: MAP2 (pink) neuronal marker. Right: quantification of Ca_V_1.2 cluster size, cluster density, and mean gray value (MGV) in the soma (a.) and dendrites (b.) of CTL (black) and PFF-treated (red) neurons. Dendritic measurements are shown separately for excitatory (dark blue) and inhibitory (light blue) populations. n = 20 somata per condition; n = 20 dendrites per group (CTL excitatory, CTL inhibitory, PFF excitatory, PFF inhibitory); two independent isolations with each isolation containing 8-10 pups. **C** Left: representative super-resolution TIRF localization maps showing Ca_V_1.2 immunolabeling in CTL (black) and PFF-treated (red) neurons. Right: quantification of PM Ca_V_1.2 cluster size, cluster density, and nearest-neighbor distance in the somatic region. n = 16 neurons per condition; two independent isolations. **D** Same experimental design as in (**B**), with neurons immunolabeled for Ca_V_2.1. n = 19 (CTL) and n = 20 (PFF) somata; n = 20 dendrites per group (CTL excitatory, CTL inhibitory, PFF excitatory, PFF inhibitory); two independent isolations. **E** Same experimental design as in (**C**), with neurons immunolabeled for Ca_V_2.1. n = 16 neurons per condition; two independent isolations. Error bars represent SEM. Statistical significance was determined using two-tailed Mann-Whitney or unpaired two-tailed t-tests. ns, not significant; *P ≤ 0.05; ***P ≤ 0.001; ****P ≤ 0.0001. CTL, control; PFF, α-synuclein pre-formed fibril treatment.

To determine whether PFF-induced remodeling extends beyond L-type channels, we next examined the distribution of additional Ca_V_ subtypes expressed in cortical neurons. We immunolabeled neurons for the P/Q-type calcium channel Ca_V_2.1^59^ and examined its distribution using the same imaging approaches. PFF treatment led to a significant increase in Ca_V_2.1 clustering in both somatic and dendritic regions (**Fig. 1D, E**), revealing a remodeling pattern similar to that observed for Ca_V_1.2 (**Fig. 1B, C**). Notably, treatment with Type II fibrils, which have a lower capacity to seed endogenous α-synuclein aggregation^60^, failed to remodel Ca_V_1.2 clusters (**Fig. S1E**), underscoring the functional distinction between Type I and Type II PFFs. Collectively, these data demonstrate that PFF exposure triggers the redistribution of multiple Ca_V_ subtypes at the neuronal plasma membrane.

### PFF treatment induces phosphorylation-dependent remodeling of the K_V_2.1 channel scaffold

Clustered K_V_2.1 channels function as structural organizers of neuronal Ca_V_ channels to shape local Ca²⁺ signaling ^32,61^. Given that PFF treatment enhanced Ca_V_ clustering at the PM (**Fig. 1**), we reasoned that this redistribution may result from PFF-dependent modifications of K_V_2.1 organization following PFF exposure. We first assessed total K_V_2.1 expression levels by WB in response to PFF treatment and observed no significant change (**Fig. S1F**).

Airyscan confocal imaging of immunolabeled K_V_2.1 revealed a modest yet significant increase in cluster density upon PFF treatment, whereas cluster size and MGV remained unchanged in both excitatory and inhibitory neurons (**Fig. 2A**). To resolve finer spatial details, we generated localization maps from super-resolution single-molecule TIRF microscopy (**Fig. 2B**). These maps corroborated the Airyscan findings (**Fig. 2A**) and further revealed that the larger clusters resolved by Airyscan consist of discrete K_V_2.1 subclusters and/or individual channels (**Fig. 2B**). Nearest neighbor distance analysis showed that PFF treatment decreased the spacing between these subclusters (**Fig. 2B**), a change that fell below the effective resolution of Airyscan imaging (**Fig. 2A**), providing evidence for PFF-dependent nanoscale reorganization of K_V_2.1.

**Fig. 2.**
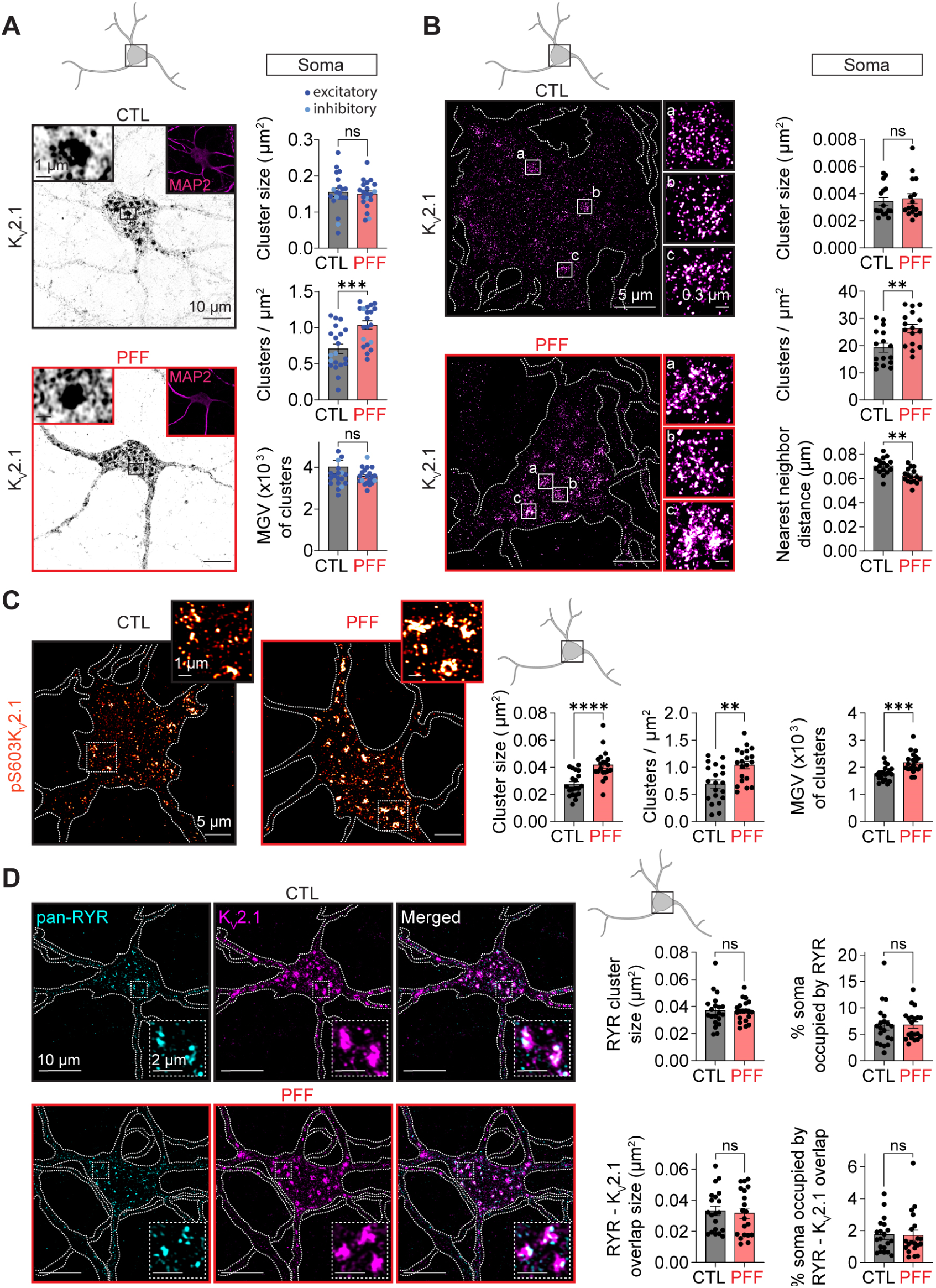
PFF treatment induces phosphorylation-dependent remodeling of K_V_2.1 at the plasma membrane. **A** Left: representative single-plane Airyscan confocal images of the PM showing K_V_2.1 immunolabeling in control (CTL, black) and PFF-treated (red) neurons. Inset: MAP2 (pink) neuronal marker. Right: quantification of K_V_2.1 cluster size, cluster density, and mean gray value (MGV) in the somatic region of CTL (black) and PFF-treated (red) excitatory (dark blue) and inhibitory (light blue) neurons. n = 20 somata per condition; two independent isolations. **B** Left: representative super-resolution TIRF localization maps showing K_V_2.1 immunolabeling in CTL (black) and PFF-treated (red) neurons. Right: quantification of PM K_V_2.1 cluster size, cluster density, and nearest-neighbor distance in the somatic region. n = 16 neurons per condition; two independent isolations. **C** Left: representative FV4000 confocal images showing pS603-K_V_2.1 immunolabeling at the PM in CTL (black) and PFF-treated (red) neurons. Images are maximum intensity projections from Z-stacks spanning whole cells. Right: quantification of PM pS603-K_V_2.1 cluster size, cluster density, and MGV in the somatic region. n = 20 neurons per condition; two independent isolations. **D** Left: representative FV4000 confocal images showing co-immunolabeling of RyR and K_V_2.1 at the PM in control (CTL, black) and PFF-treated (red) neurons. Images are maximum intensity projections from ten optical sections acquired at the PM. Right: quantification of RyR cluster size, RyR-K_V_2.1 overlap area, and somatic occupancy (% of soma area) for RyR clusters and RyR-K_V_2.1 overlap in CTL (black) and PFF-treated (red) neurons. n = 20 neurons per condition; two independent isolations. Error bars represent SEM. Statistical significance was determined using two-tailed Mann-Whitney or unpaired two-tailed t-tests. ns, not significant; ns, not significant, **P ≤ 0.01; ***P ≤ 0.001; ****P ≤ 0.0001. CTL, control; PFF, α-synuclein pre-formed fibril treatment; pS603-K_V_2.1, K_V_2.1 phosphorylated at serine 603.

K_V_2.1 clustering is dynamically regulated by its phosphorylation state: kinase-mediated phosphorylation promotes clustering, whereas phosphatase-driven dephosphorylation induces cluster dispersal^35,36,62^. To determine whether the increased subcluster proximity was associated with altered phosphorylation, we immunolabeled neurons for the S603 phosphorylation site on K_V_2.1 (pS603-K_V_2.1), a modification known to promote channel clustering^35,36,63^. Confocal imaging revealed that phosphorylation at this residue was significantly increased following PFF treatment, with pS603-K_V_2.1 exhibiting increased cluster size, density, and MGV (**Fig. 2C**). These data indicate that elevated S603 phosphorylation promotes tighter association of K_V_2.1 subclusters, accounting for the reduced inter-subcluster spacing observed by TIRF (**Fig. 2B**).

In addition to Ca_V_ channels, K_V_2.1 also have the capacity to organize ER-localized RyRs to shape local Ca^2+^ signals ^64^. We therefore investigated whether PFF treatment remodels RyR distribution by simultaneously labeling K_V_2.1 at the PM and RyR at the ER. Multiplexed immunolabeling followed by confocal imaging revealed that RyR distribution and the overall organization between K_V_2.1-RyR clusters, remained preserved following PFF treatment (**Fig. 2D**). Together, these findings suggest that PFF-dependent alterations in K_V_2.1 phosphorylation remodel the channel scaffold at the PM without disrupting broader interactions with nearby RyR.

### PFF exposure increases the spatial proximity of Ca_V_ and K_V_2.1 channels

Given that PFF treatment enhances Ca_V_ clustering (**Fig. 1**) and that K_V_2.1 plays an established role in organizing Ca_V_ distribution^32,65^, we next examined whether PFF exposure alters their colocalization and whether disrupting K_V_2.1–Ca_V_1.2 coupling influences the PFF-induced changes in Ca_V_ clustering. To test this, we used a synthetic, membrane-permeable peptide corresponding to the <u>c</u>alcium <u>c</u>hannel <u>a</u>ssociation <u>d</u>omain (CCAD) in the cytoplasmic C-terminus of K_V_2.1 ^32^. This peptide mimics the CCAD region and acts as a competitive inhibitor that selectively disrupts Ca_V_-K_V_2.1 coupling (**Fig. 3A**) ^28,32^. A scrambled peptide (SCRBL) served as control. Neurons were incubated with CCAD or SCRBL peptides before immunolabeling for K_V_2.1 and Ca_V_1.2 and subsequent Airyscan imaging. Quantitative analysis revealed that PFF treatment significantly increased both Ca_V_1.2 cluster size and the extent of Ca_V_1.2-K_V_2.1 overlap at the somatic PM (**Fig. 3B**). The CCAD peptide not only disrupted Ca_V_1.2-K_V_2.1 coupling but also restored Ca_V_1.2 cluster size to control levels, indicating that PFF-induced increases in Ca_V_1.2 clustering are driven by alterations in K_V_2.1 scaffolding.

**Fig. 3.**
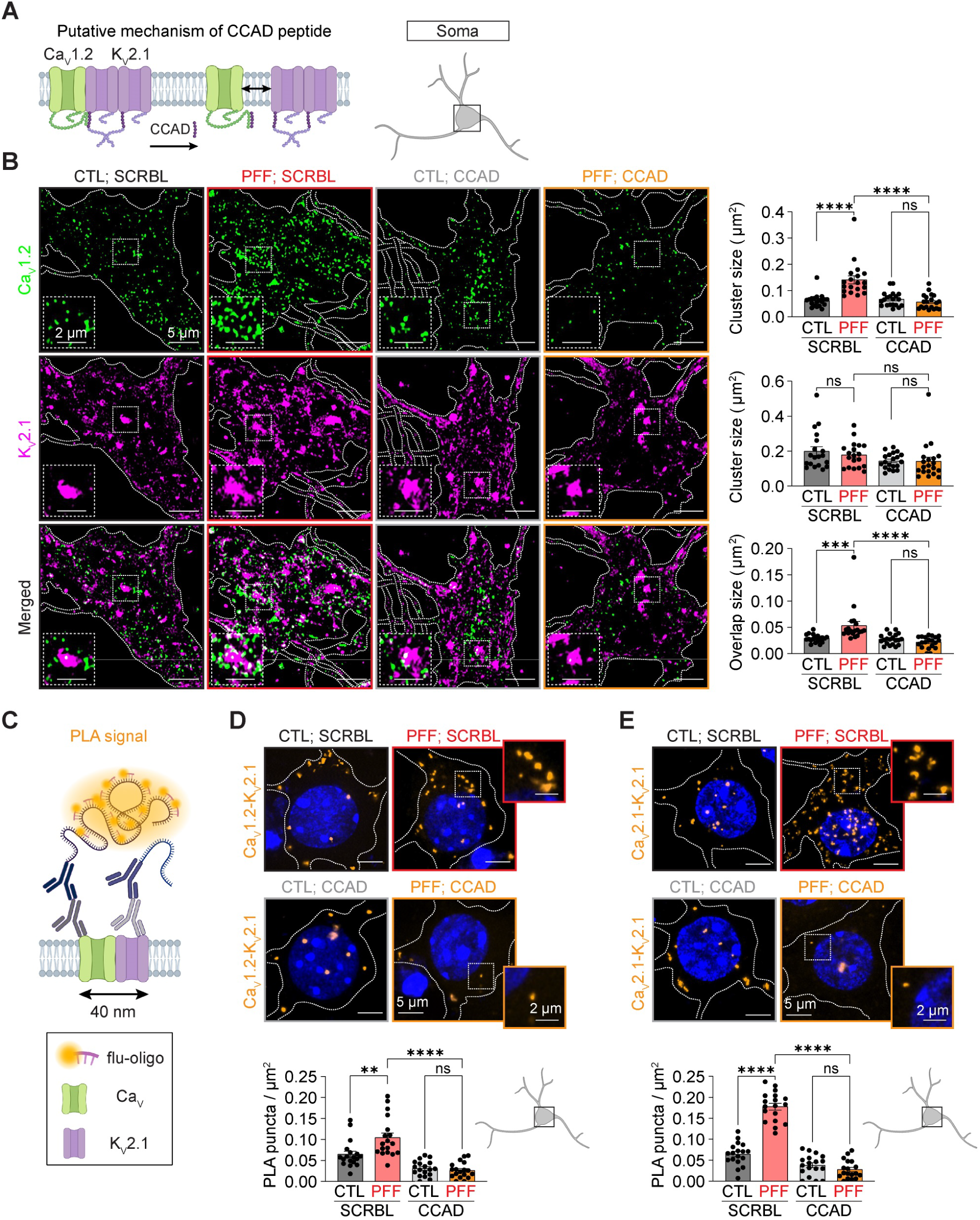
PFF treatment increases Ca_V_-K_V_2.1 spatial proximity in a K_V_2.1 scaffolding-dependent manner. **A** Schematic representation of the CCAD peptide mechanism of action. **B** Left: representative single-plane Airyscan confocal images of the PM in CTL and PFF-treated neurons co-incubated with SCRBL or CCAD peptides and co-immunolabeled for Ca_V_1.2 and K_V_2.1. Conditions are shown as CTL;SCRBL (black), PFF;SCRBL (red), CTL;CCAD (gray), and PFF;CCAD (yellow). Right: quantification of Ca_V_1.2 cluster size, K_V_2.1 cluster size, and Ca_V_1.2-K_V_2.1 overlap area in the somatic region. n = 19 (CTL;SCRBL), n = 19 (PFF;SCRBL), n = 20 (CTL;CCAD), and n = 20 (PFF;CCAD) neurons; two independent isolations. **C** Schematic representation of the proximity ligation assay (PLA). **D** Top: representative Airyscan confocal PLA images showing Ca_V_1.2-K_V_2.1 proximity. Images are maximum intensity projections from Z-stacks spanning whole cells. Color coding as in (**B**). Bottom: quantification of PLA puncta density. n = 18 neurons per condition; two independent isolations. **E** Same experimental design as in (**D**), but assessing Ca_V_2.1-K_V_2.1 proximity. n = 18 neurons per condition; two independent isolations. Error bars represent SEM. Statistical significance in panels (**B, D-E**) was determined using two-way ANOVA with appropriate post hoc tests. ns, not significant; **P ≤ 0.01; ***P ≤ 0.001; ****P ≤ 0.0001. CTL, control; PFF, α-synuclein pre-formed fibril treatment; CCAD, calcium channel association domain peptide; SCRBL, scrambled control peptide.

To independently assess channel proximity at higher resolution, we performed proximity ligation assay (PLA), which generates a discrete fluorescent signal when two target proteins are within approximately 40 nm (**Fig. 3C**) (see Methods)^28^. PLA analysis revealed a significant increase in Ca_V_1.2-K_V_2.1 proximity following PFF treatment, reflected by increased puncta density compared to controls (**Fig. 3D**). This increase was effectively reversed by the CCAD peptide (**Fig. 3D**). PLA experiments assessing Ca_V_2.1-K_V_2.1 proximity yielded similar results (**Fig. 3E**), demonstrating that the PFF-dependent increase in Ca_V_-K_V_2.1 association extends across Ca_V_ subtypes. Control experiments confirmed that the CCAD peptide did not alter K_V_2.1 clustering itself, and assay specificity was verified by omitting one of the primary antibodies (**Fig. S2**). Together, these data demonstrate that PFF treatment increases Ca_V_-K_V_2.1 nanoscale proximity and that PFF-dependent Ca_V_ clustering relies on the scaffolding function of K_V_2.1.

### CDK5 inhibition abolishes PFF-induced K_V_2.1-Ca_V_ proximity

Cyclin-dependent kinase 5 (CDK5) is a key neuronal kinase that phosphorylates K_V_2.1 at S603 within the C-terminal phosphoregulatory domain to promote channel clustering^35^. CDK5 activity is elevated in multiple models of PD, where extracellular α-synuclein has been shown to induce calpain-dependent CDK5 overactivation, and CDK5 hyperactivity has been identified as a mediator of dopaminergic neuron loss ^66,67^. Given that PFF exposure increased S603 phosphorylation (**Fig. 2C**), we asked whether CDK5 contributes to this effect. Co-immunolabeling of CDK5 and K_V_2.1 followed by confocal imaging at the PM revealed a significant increase in their spatial overlap following PFF treatment, whereas CDK5 puncta density remained unchanged (**Fig. 4A**). Analysis of whole-cell cross-sections confirmed similar subcellular distribution and overall expression levels of CDK5 in control and PFF-treated neurons (**Fig. S3A**), suggesting that PFF exposure enhances the spatial association of CDK5 with K_V_2.1 rather than altering CDK5 expression or global distribution.

**Fig. 4.**
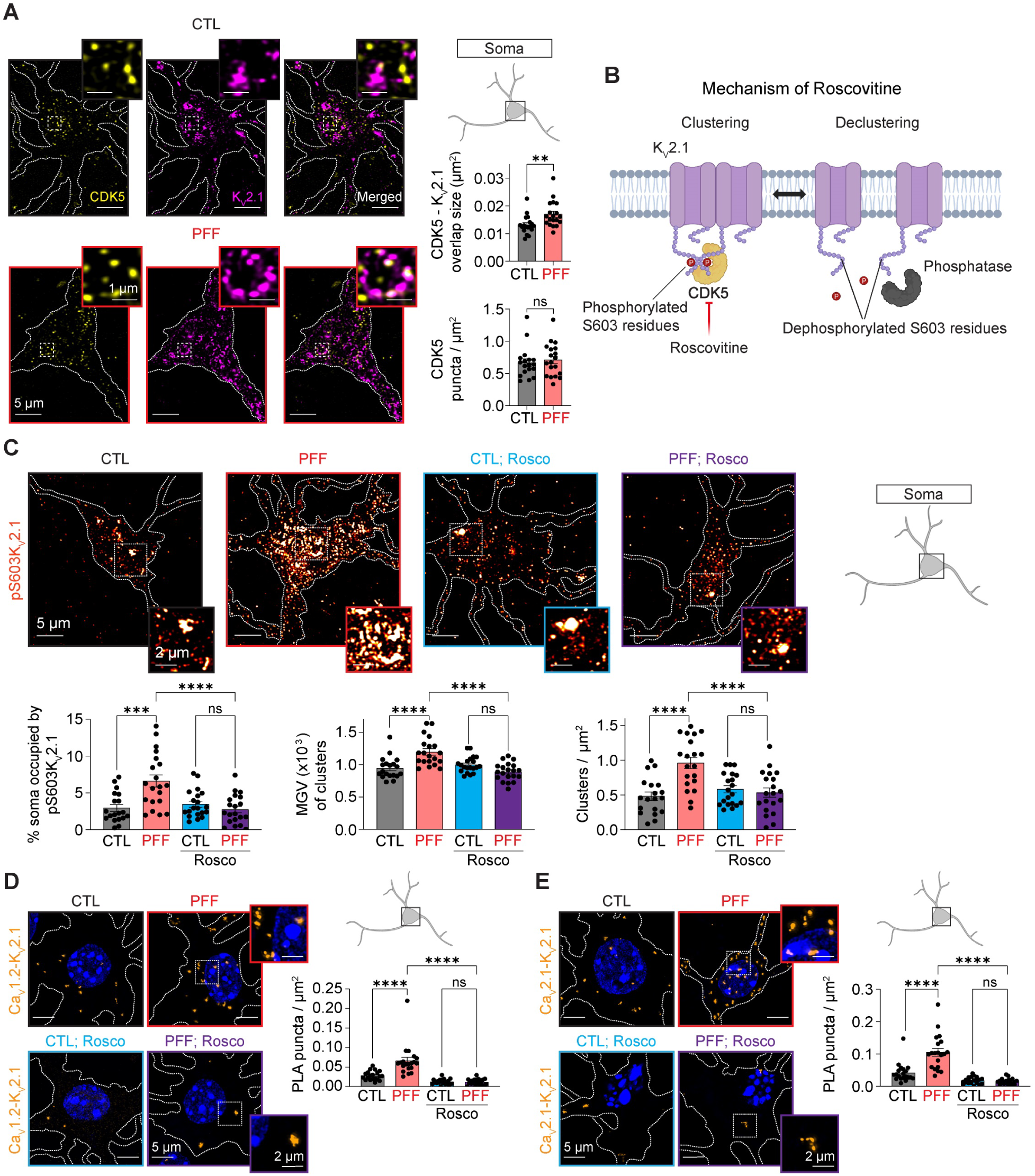
CDK5 inhibition attenuates PFF-dependent K_V_2.1 phosphorylation and Ca_V_-K_V_2.1 proximity. **A** Left: representative FV4000 confocal images showing co-immunolabeling of CDK5 and K_V_2.1 at the PM in control (CTL, black) and PFF-treated (red) neurons. Images are maximum intensity projections from three optical sections acquired at the PM. Right: quantification of CDK5-K_V_2.1 overlap area and CDK5 puncta density in the somatic region. n = 18 (CTL) and n = 19 (PFF) neurons; two independent isolations. **B** Schematic representation of the roscovitine mechanism of action. **C** Top: representative FV4000 confocal images of the PM in CTL and PFF-treated neurons incubated with or without roscovitine and immunolabeled for pS603-K_V_2.1. Conditions are shown as CTL (black), PFF (red), CTL;Rosco (blue), and PFF;Rosco (purple). Images are maximum intensity projections from Z-stacks spanning whole cells. Bottom: quantification of somatic pS603-Kv2.1 occupancy (% of soma area), cluster MGV, and cluster density. n = 19 (CTL), n = 20 (PFF), n = 20 (CTL;Rosco), and n = 20 (PFF;Rosco) neurons; two independent isolations. **D** Left: representative FV4000 confocal PLA images showing Ca_V_1.2-K_V_2.1 proximity. Images are maximum intensity projections from Z-stacks spanning whole cells. Color coding as in (**C**). Right: quantification of PLA puncta density. n = 20 neurons per condition; two independent isolations. **E** Same experimental design as in (**D**), but assessing Ca_V_2.1-K_V_2.1 proximity. n = 20 (CTL), n = 20 (PFF), n = 21 (CTL;Rosco), and n = 20 (PFF;Rosco) neurons; two independent isolations. Error bars represent SEM. Statistical significance in panel (**A**) was determined using two-tailed Mann-Whitney test; panels (**C**–**E**) were analyzed using two-way ANOVA with appropriate post hoc tests. ns, not significant; **P ≤ 0.01; ***P ≤ 0.001; ****P ≤ 0.0001. CTL, control; PFF, α-synuclein pre-formed fibril treatment; Rosco, roscovitine; pS603-K_V_2.1, K_V_2.1 phosphorylated at serine 603.

To directly test the contribution of CDK5 to PFF-induced K_V_2.1 phosphorylation, we inhibited CDK5 with roscovitine (10 μM) (**Fig. 4B**) and assessed the phosphorylation state of K_V_2.1 by immunolabeling for pS603-K_V_2.1. CDK5 inhibition restored PFF-dependent K_V_2.1 phosphorylation to control levels, as reflected by reduced pS603-K_V_2.1 occupancy in the soma, cluster MGV, and cluster density (**Fig. 4C).** Notably, pS603-K_V_2.1 levels in the control group were unaltered by roscovitine treatment, which we attribute to the reduced network activity typical of cultured neurons, where basal Ca^2+^-dependent phosphatase activity such as calcineurin may remain low, limiting K_V_2.1 dephosphorylation even when CDK5 is inhibited^68^. We next asked whether reducing K_V_2.1 phosphorylation through CDK5 inhibition alters Ca_V_-K_V_2.1 association. PLA analysis confirmed that PFF treatment increased Ca_V_1.2-K_V_2.1 proximity (consistent with **Fig. 3D**), while roscovitine restored PLA puncta density to control levels (**Fig. 4D**). PLA experiments assessing Ca_V_2.1-K_V_2.1 proximity yielded similar results (**Fig. 4E**), and negative controls verified assay specificity (**Fig. S3B**). Collectively, these findings indicate that PFF treatment enhances CDK5-dependent phosphorylation of K_V_2.1, which in turn promotes Ca_V_ channel retention and clustering at K_V_2.1-organized sites.

### PFF-dependent Ca_V_ clustering elevates Ca^2+^ influx and drives aberrant c-Fos expression

Ca_V_ channel clustering increases channel open probability and promotes cooperative gating to enhance Ca^2+^ influx^23^. Our finding that PFF treatment enhances Ca_V_ clustering at K_V_2.1-organized sites (**Figs. 1,3**) prompted us to determine whether this remodeling is accompanied by elevated Ca^2+^ influx in this PD model (**Fig. 5A**).

**Fig. 5.**
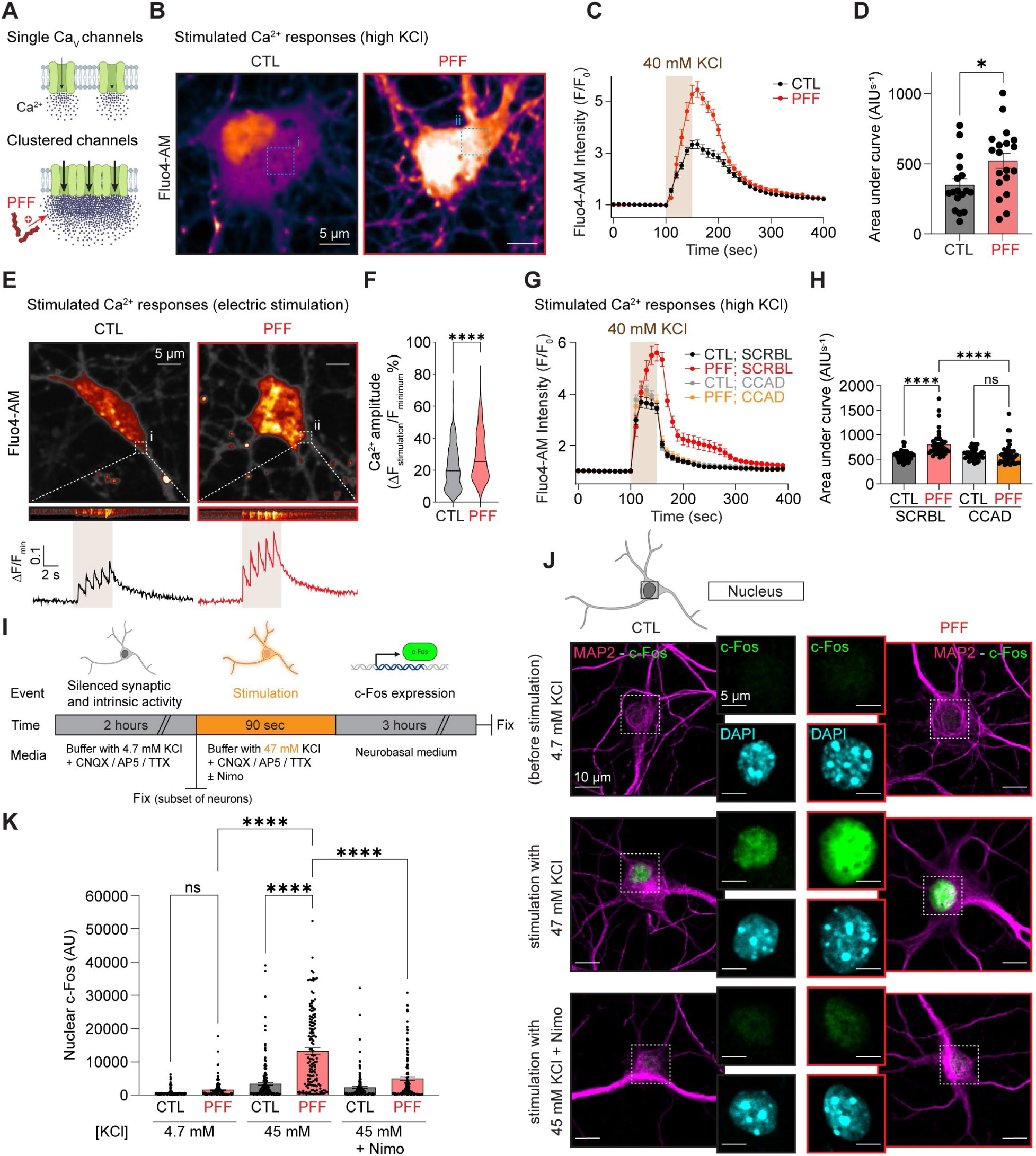
PFF-dependent Ca_V_ clustering elevates depolarization-evoked Ca^2+^ influx and drives aberrant c-Fos expression. **A** Schematic of the proposed model. **B** Representative spinning-disk confocal images of Fluo-4-loaded CTL (black) and PFF-treated (red) neurons stimulated with 40 mM KCl. Cytoplasmic ROIs (i–ii) were analyzed. **C** Averaged and normalized Fluo-4 fluorescence time series from experiment (**B**). **D** Quantified area under curve from normalized Fluo-4 intensity traces. Points represent averaged ROIs from a single field of view. n = 170 (CTL) and n = 86 (PFF) neurons; three independent isolations. **E** Top: Same experimental design as in (**B**), except neurons were stimulated with electrical field stimulator. Bottom: kymographs and normalized Fluo-4 fluorescence time series from ROIs (i–ii). **F** Quantified Ca^2+^ peak amplitude upon electrical field stimulation. n = 490 (CTL) and n = 560 (PFF) peaks; two independent isolations. **G** Same experimental design as in (**C**), except neurons were co-incubated with SCRBL or CCAD peptides. Whole-cell somata were analyzed. **H** Quantified area under curve from normalized Fluo-4 intensity traces. n = 52 (CTL;SCRBL), n = 46 (PFF;SCRBL), n = 47 (CTL;CCAD), and n = 50 (PFF;CCAD) neurons; two independent isolations. **I** Experimental design for c-Fos induction. **J** Representative single-plane FV4000 confocal images at nuclear cross-sections showing c-Fos immunolabeling in CTL (black) and PFF-treated (red) neurons under basal conditions (4.7 mM KCl), depolarization (47 mM KCl), or depolarization with Ca_V_1 inhibition (47 mM KCl + nimodipine). MAP2 marks neurons; DAPI labels nuclei. **K** Quantified nuclear c-Fos immunoreactivity in CTL (black) and PFF-treated (red) neurons under the conditions indicated. n = 136 (CTL;4.7 mM KCl), n = 210 (CTL;47 mM KCl), n = 156 (CTL;47 mM KCl + Nimo), n = 113 (PFF;4.7 mM KCl), n = 145 (PFF;47 mM KCl), and n = 147 (PFF;47 mM KCl + Nimo) neurons; two independent isolations. Error bars represent SEM. Statistical significance: unpaired two-tailed t-test (**D**), two-tailed Mann-Whitney test (**F**), or two-way ANOVA (**H**, **K**). ns, not significant; *P ≤ 0.05; ****P ≤ 0.0001. CTL, control; PFF, α-synuclein pre-formed fibril treatment; CCAD, calcium channel association domain peptide; SCRBL, scrambled control peptide; TTX, tetrodotoxin; Nimo, nimodipine.

To quantify activity-evoked Ca^2+^ entry, before confocal microscopy neurons were loaded with the membrane-permeable Ca²⁺ indicator Fluo-4 AM, which increases fluorescence upon binding free cytosolic Ca^2+ 69^. Without stimulation, both control and PFF-treated neurons showed similar cytosolic Ca²⁺ levels, as reflected by the MGV of Fluo-4 fluorescence (**Fig. S4A**). However, depolarization with 40 mM KCl, which promotes Ca²⁺ influx through Ca_V_ channels^70^, elicited significantly larger Ca^2+^ transients in PFF-treated neurons, reflected by both increased peak fluorescence and a greater integrated response (**Fig. 5B–D**). Electrical field stimulation produced similarly elevated Ca^2+^ amplitudes in PFF-treated neurons, confirming that the enhanced response persists under a more physiological activation paradigm (**Fig. 5E, F**).

To test whether the elevated Ca^2+^ influx depends on Ca_V_-K_V_2.1 coupling, we applied the CCAD peptide during KCl-evoked depolarization. CCAD treatment reduced the PFF-dependent Ca^2+^ increase to control levels (**Fig. 5G, H**), demonstrating that K_V_2.1-mediated recruitment of Ca_V_ channels is required for the enhanced Ca²⁺ response. Beyond elevated cytosolic Ca^2+^, PFF-treated neurons also exhibited increased nuclear Fluo-4 fluorescence upon stimulation (**Fig. S4B**), raising the possibility that enhanced Ca^2+^ entry may influence nuclear signaling and gene expression.

Ca_V_ channels are established mediators of excitation-transcription coupling, linking activity-dependent Ca^2+^ influx to signaling pathways that regulate gene expression ^71^. Given that PFF treatment amplifies both cytosolic and nuclear Ca^2+^ signals (**Fig. 5B–H; Fig. S4B**), and that transcriptomic studies have identified widespread gene expression changes in PD^40^, we asked whether α-synuclein pathology also drives aberrant Ca^2+^-dependent transcriptional responses. We focused on c-Fos, an immediate early gene and Ca²⁺-responsive transcription factor whose expression is tightly regulated by neuronal activity and Ca²⁺-dependent signaling^38,72^.

To assess activity-dependent c-Fos induction, neurons were first silenced by incubation with CNQX, AP5, and tetrodotoxin (TTX) to block AMPA/NMDA receptors and voltage-gated Na⁺ channels, respectively, thereby isolating Ca_V_-dependent depolarization responses (**Fig. 5I**). Neurons were then depolarized with KCl (final concentration 47 mM) in the continued presence of these blockers and returned to conditioned medium to permit c-Fos expression (**Fig. 5I**; see Methods). Baseline nuclear c-Fos immunoreactivity was similar between control and PFF-treated groups (**Fig. 5J, K**). However, depolarization evoked a significantly greater c-Fos response in PFF-treated neurons (**Fig. 5J, K**), consistent with the enhanced cytosolic and nuclear Ca^2+^ signals observed upon stimulation (**Fig. 5B–H; Fig. S4B**).

To determine whether Ca_V_ channel activity drives this transcriptional amplification, we inhibited L-type Ca_V_ channels with nimodipine during depolarization. Nimodipine significantly attenuated the PFF-dependent increase in c-Fos expression (**Fig. 5J, K**), indicating that Ca_V_-mediated Ca²⁺ entry is a critical contributor to the aberrant excitation-transcription coupling observed in PFF-treated neurons.

Together, these findings demonstrate that PFF-driven remodeling of Ca_V_ channel nanodomains amplifies stimulus-evoked Ca²⁺ influx, which in turn drives aberrant Ca^2+^-dependent gene expression, establishing a direct functional link between α-synuclein-induced channel reorganization and transcriptional dysregulation.

## Discussion

In this study, we identify a mechanism by which α-synuclein pathology drives excessive Ca^2+^ influx at the neuronal plasma membrane and demonstrate how this aberrant Ca^2+^ entry disrupts excitation-transcription coupling. Using a model of idiopathic PD, we show that α-synuclein PFFs markedly increase Ca_V_ channel clustering at the PM, where K_V_2.1 channels act as critical scaffolds that organize Ca_V_ nanodomains at ER-PM junctions. This remodeling is driven by CDK5-dependent hyperphosphorylation of K_V_2.1, which enhances its scaffolding capacity and promotes Ca_V_ retention. Consequently, depolarization evokes exaggerated Ca^2+^ entry into the cytoplasm, amplifying Ca^2+^-dependent gene expression programs. Importantly, disrupting Ca_V_–K_V_2.1 coupling significantly reduced Ca²⁺ influx, while pharmacological inhibition of Ca_V_ channels significantly attenuated the PFF-induced changes in gene expression. Together, our data reveal that α-synuclein-driven nanoscale reorganization of Ca_V_ channels reinforces maladaptive transcriptional responses, providing a mechanistic link between altered Ca^2+^ signaling, aberrant gene expression, and neuronal dysfunction in PD (**Fig. 6**).

**Fig. 6.**
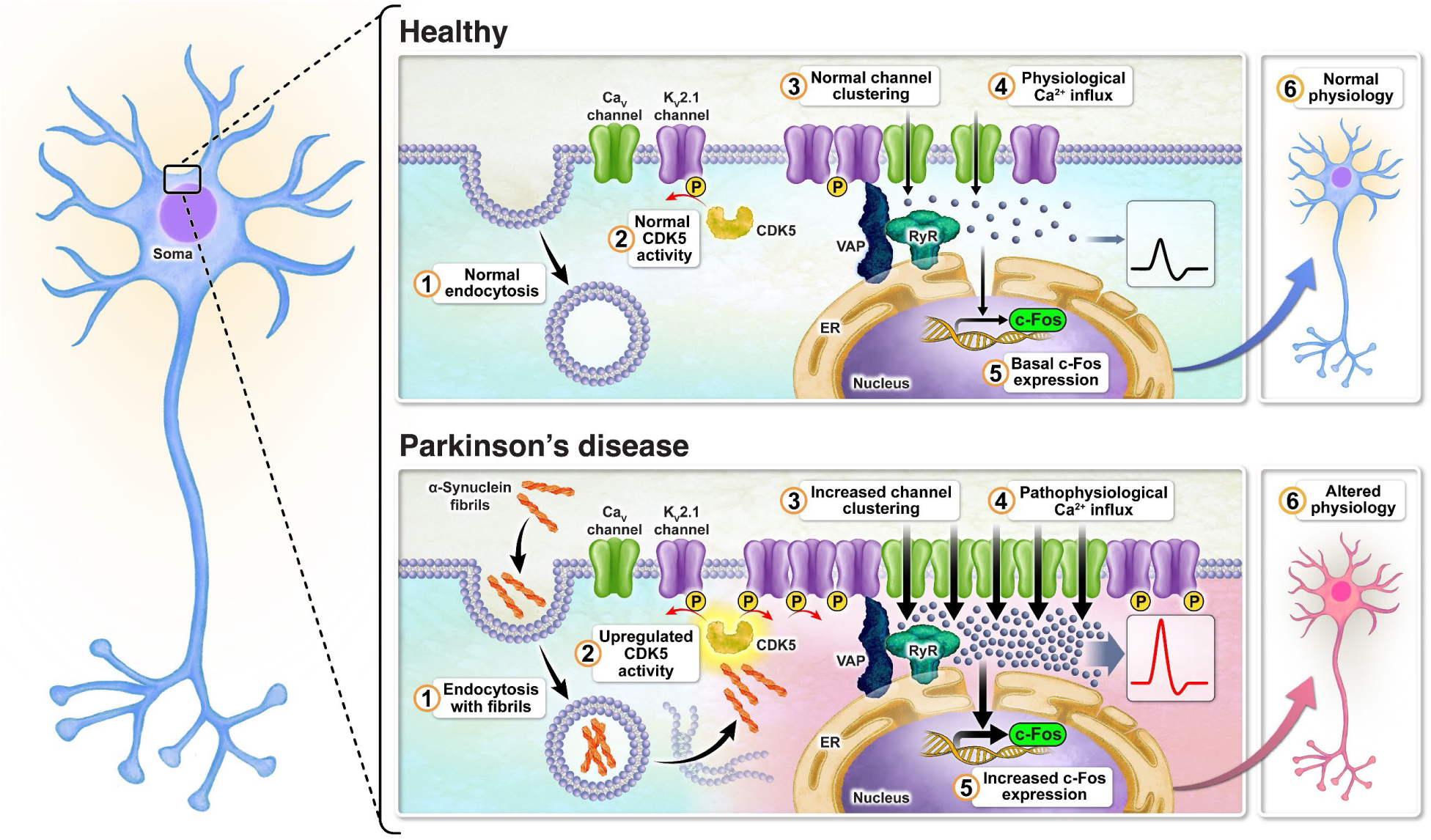
model: α-synuclein fibrils drive the remodeling of voltage-gated cation channels, increase Ca^2+^ influx, and promote transcriptional dysregulation.

PD is a slowly progressing disorder characterized by the accumulation of α-synuclein aggregates that form Lewy bodies, leading to widespread cellular dysfunction and progressive neurodegeneration^73–75^. A critical hallmark beyond Lewy bodies is disrupted Ca^2+^ homeostasis ^18,51,76,77^. Although it is well established that Ca^2+^ transfer from the ER lumen to mitochondria can result in excessive mitochondrial Ca^2+^ uptake and elevated oxidative stress^18,78–80^, the mechanism by which Ca^2+^ is replenished from the extracellular space has remained less clear, despite neurons relying on extracellular Ca^2+^ as their ultimate source. Voltage-gated Ca^2+^ channels represent strong candidates in this process, as they mediate the primary depolarization-evoked Ca^2+^ entry and their gating properties are amenable to regulatory modulation^23^. Indeed, we find that PFF exposure induces a reorganization of Ca_V_ channels at the neuronal PM, leading to enhanced clustering and increased Ca^2+^ influx upon depolarization. This is consistent with the biophysical consequences of channel clustering, as nanoscale proximity between Ca_V_ channels can promote functional coupling, enhance cooperative gating, and increase local open probability, thereby amplifying macroscopic Ca^2+^ current density^20,22^. Our findings add to a growing body of evidence that Ca_V_ channel clustering is altered in disease contexts, aging and type 2 diabetes, where disrupted nanoscale channel organization contributes to cellular dysfunction^52,53^.

We find that only Type I PFFs induce increased Ca_V_ clustering, whereas Type II PFFs fail to remodel Ca_V_1.2 clusters. This suggests that the higher seeding efficiency of Type I PFFs and their enhanced capacity to convert endogenous α-synuclein monomers into aggregated species^60^ are required for Ca_V_ remodeling. Consistent with this interpretation, we observe increased Ca_V_ clustering in inhibitory neurons, despite their low endogenous α-synuclein expression and relative resistance to Lewy body formation^47,81^, indicating that Type I PFFs seed endogenous α-synuclein aggregation within these neurons as well. This aligns with the emerging view that smaller oligomeric α-synuclein species, rather than large Lewy body inclusions, represent the primary toxic forms responsible for neurodegeneration ^82,83^.

We focused our analysis on the neuronal soma because this compartment harbors the highest density of K_V_2.1-organized ER-PM junctions and their associated Ca_V_ clusters^32,65^, and because the soma is the primary site of excitation-transcription coupling, where Ca^2+^ signals generated at the PM are decoded by perinuclear and nuclear signaling machinery to regulate gene expression^84^. Our imaging data confirm that Ca_V_ channels are preferentially enriched at K_V_2.1-organized somatic domains, where K_V_2.1 serves as a structural scaffold that stabilizes Ca_V_ clusters. This organization positions somatic Ca_V_-K_V_2.1 complexes as the most functionally relevant targets for understanding how PFF-dependent channel remodeling impacts transcriptional output. Consistent with this, treatment with the CCAD peptide disrupts Ca_V_-K_V_2.1 spatial coupling, leading to Ca_V_ declustering, reduced nanoscale proximity, and a consequent reduction in depolarization-evoked Ca^2+^ influx.

Within this regulatory framework, we identify S603 on K_V_2.1^35^ as being robustly phosphorylated in a PFF-dependent manner by CDK5. Pharmacological inhibition of CDK5 with roscovitine reduces the nanoscale proximity between Ca_V_ and K_V_2.1 channels. These findings support a model in which phosphorylation at S603 enhances the scaffolding function of K_V_2.1, promoting Ca_V_ clustering and facilitating Ca_V_ retention at ER-PM junctions. The precise molecular basis of this co-clustering, whether through direct protein-protein interaction or scaffold-mediated mechanisms, remains to be established.

Our findings are consistent with several studies demonstrating that CDK5 activity is elevated in PD ^66,67,85,86^. The best characterized mechanism involves calpain-mediated cleavage of the membrane-bound p35/CDK5 complex into the cytoplasmic p25/CDK5 complex, resulting in spatially deregulated phosphorylation of downstream substrates ^67,86^. This appears paradoxical for K_V_2.1, which is a PM-localized substrate^87^. We propose that an alternative pathway may be responsible: CDK5 activity can be enhanced by phosphorylation at Tyr15 ^67,88^, a modification mediated by upstream kinases such as c-Abl^89^, which is activated in response to α-synuclein-induced oxidative stress^90,91^. This regulatory mechanism would allow CDK5 hyperactivation at the PM without requiring p25-dependent mislocalization and is consistent with the increased CDK5-K_V_2.1 spatial overlap we observe without changes in overall CDK5 distribution. Further work will be needed to distinguish these pathways and determine their relative contributions to K_V_2.1 hyperphosphorylation in disease.

A related question concerns the role of calcineurin, which would be expected to dephosphorylate K_V_2.1 in response to elevated Ca^2+^ levels, promoting cluster dispersal and restoring channel conductance^36^. We propose that sustained CDK5 activity outpaces the periodic calcineurin-mediated dephosphorylation, the net result would be hyperphosphorylation of K_V_2.1, maintaining Ca_V_ channels in a clustered, high-activity configuration. Testing this kinase-phosphatase balance model will be an important direction for future studies.

Among second messengers, Ca^2+^ uniquely mediates excitation-transcription coupling in neurons by activating Ca^2+^-dependent signaling pathways that regulate gene expression^38,71^, with Ca_V_ channels serving as the principal source of activity-dependent Ca^2+^ influx^71^. Ca^2+^ homeostasis is disrupted in PD and associated with altered gene expression patterns ^40^. Consistent with this, we observe a PFF-dependent increase in depolarization-induced c-Fos expression, an effect attenuated by pharmacological inhibition of Ca_V_ channels. This underscores the importance of PFF-induced Ca_V_ remodeling and clustering in driving aberrant excitation-transcription coupling.

The downstream consequences of elevated c-Fos expression may be particularly significant in the context of neurodegeneration. Under physiological conditions, c-Fos plays a central role in cognitive processes, including learning and memory, by regulating gene programs that underlie synaptic plasticity and long-term neuronal adaptations^92,93^. However, sustained or excessive c-Fos activity can become maladaptive. Elevated c-Fos expression is reported in multiple studies of Alzheimer’s disease, where it is associated with cognitive decline and increased neuronal apoptosis^94,95^. Similarly, increased c-Fos expression is observed across multiple brain regions in various PD models, including cortical areas and the hippocampus^96–98^. c-Fos, as a component of the AP-1 transcription factor complex, regulates the expression of pro-apoptotic genes such as Bim and FasL, and sustained AP-1 activation in the context of chronic cellular stress can shift the transcriptional balance from pro-survival to pro-death programs^99^. In PFF-treated neurons, where Ca_V_-mediated Ca^2+^ influx is amplified with each depolarization cycle, the repeated induction of c-Fos and its downstream targets may progressively overwhelm neuronal stress responses, contributing to a transcriptional state that favors degeneration over adaptation. Elucidating the specific signaling cascades linking Ca_V_-dependent Ca^2+^ entry to c-Fos induction, and identifying the broader gene expression programs activated downstream, will be critical for understanding how Ca^2+^-dependent transcriptional dysregulation contributes to neuronal vulnerability in PD.

Our finding that PFF-treated neurons exhibit higher-amplitude Ca^2+^ transients upon stimulation, resulting in amplified activation of Ca^2+^-dependent signaling pathways, has implications beyond excitation-transcription coupling. Elevated Ca^2+^ influx through remodeled Ca_V_ clusters is expected to impact multiple Ca^2+^-dependent processes that are disrupted in PD. Ca^2+^ can directly bind to α-synuclein and promote its aggregation^100^, creating a feedforward loop in which Ca_V_-mediated Ca^2+^ entry accelerates the very pathology that drives channel remodeling. Excess Ca^2+^ also exacerbates mitochondrial dysfunction by enhancing Ca^2+^ transfer at ER-mitochondria contact sites, driving reactive oxygen species production and oxidative stress^18^. Furthermore, Ca^2+^ facilitates α-synuclein cell-to-cell propagation by supporting exocytotic release^101^, thereby amplifying the spread of pathology. Together, these observations suggest that Ca_V_ remodeling may serve as a central amplifier of multiple pathogenic cascades in PD, with excessive Ca^2+^ entry simultaneously promoting α-synuclein aggregation, mitochondrial dysfunction, transcriptional dysregulation, and intercellular spread of pathology.

This framework is consistent with the well-established vulnerability of substantia nigra dopaminergic neurons, which exhibit high autonomous pacemaking activity accompanied by sustained Ca^2+^ influx and express relatively low levels of the Ca^2+^-buffering protein calbindin-D28k^80^. These intrinsic properties would be expected to render dopaminergic neurons particularly sensitive to the Ca_V_ remodeling mechanism we describe, as even modest increases in Ca_V_ clustering could substantially amplify the already elevated Ca^2+^ burden associated with pacemaking^80^, accelerating downstream pathogenic processes. In the present study, we used cortical neurons, consistent with the progressive spread of α-synuclein pathology to cortical regions in later disease stages, where it drives severe cognitive decline and dementia^14,41,43,46^. Cortical neurons express α-synuclein and are susceptible to Lewy body formation^47,81^, and the robust Ca_V_-K_V_2.1 organization in these neurons provides an experimentally tractable system for dissecting the mechanisms of channel remodeling. A limitation of this approach is that cortical neurons differ from dopaminergic neurons in their intrinsic firing properties, Ca^2+^ buffering capacity, and vulnerability profile^102^. Determining whether the Ca_V_ remodeling mechanism identified here operates in dopaminergic neurons of the substantia nigra, and whether cell-type-specific differences in Ca^2+^ handling modulate its impact, will be an important direction for future investigation.

Our study identifies Ca_V_ channels as central mediators of excessive Ca^2+^ entry during α-synuclein-induced Ca^2+^ imbalance and provides a mechanistic framework in which nanoscale channel reorganization, rather than altered channel expression, drives pathological Ca^2+^ influx and aberrant gene expression. These findings position PD, at least in part, as a nanostructural channelopathy. Given the numerous intracellular roles of Ca^2+^ and the wide range of signaling cascades it controls^103^, many of which are disrupted in PD^16,103^, our data highlight Ca_V_ channels as compelling therapeutic targets. As Ca^2+^ imbalance appears to represent an early pathogenic event^103^, modulating Ca_V_ activity or disrupting Ca_V_-K_V_2.1 coupling may restore Ca^2+^ homeostasis at its source, potentially attenuating downstream pathogenic processes before irreversible neuronal damage occurs.

### Limitations and future directions

Our experiments were performed in cultured cortical neurons, a model that captures later stages of pathology when α-synuclein aggregates spread beyond the midbrain to cortical regions^46^. It will be important to test whether the same CDK5-K_V_2.1-Ca_V_ remodeling occurs in vulnerable dopaminergic populations and *in vivo*, and whether it contributes to a self-reinforcing cycle in which elevated Ca^2+^ entry further promotes α-synuclein aggregation and cellular stress. Finally, mechanisms that tune ER Ca^2+^ handling (for example, altered SERCA activity) and lipid-dependent regulation of Ca_V_ clustering (for example, phosphoinositide signaling) may intersect with this pathway and represent tractable entry points for therapeutic.

## Methods

### Animals and cell culture

All animal experiments were performed in compliance with ethical regulations approved by the University of California Davis Institutional Animal Care and Use Committee (protocol no. 22644). C57BL/6 wild-type mice were purchased from The Jackson Laboratory and housed in a vivarium under controlled conditions.

Primary cortical neurons were isolated from embryonic day 15–18 (E15–18) mice of both sexes, pooling tissue from 6–10 pups per isolation. Cortices were dissected in chilled phosphate-buffered saline (PBS) under sterile conditions. Neurons were dissociated using the Worthington Papain Dissociation System (LK003150) according to the manufacturer’s recommendations. Briefly, cortical tissue was incubated in papain solution containing DNase I for 25 min at 37 °C with gentle agitation every 5 min. Following mechanical trituration, cells were collected by centrifugation at 423 × g for 5 min and resuspended in Earle’s Balanced Salt Solution containing ovomucoid (papain inhibitor) and DNase I. Cells were centrifuged again at 423 × g for 1 min. Neurons were plated at a density of 150,000 cells/mL onto poly-D-lysine-coated (Sigma, P6407) coverslips (EMS, Platinum Line, 25 mm, No. 1.5, 71887) in six-well plates (Fisher Scientific, 08-772-1B). Culture medium consisted of 2 mL Neurobasal medium (Gibco, 21103-049) supplemented with GlutaMAX (Gibco, 35050-061), B-27 supplement (Gibco, 17504-044), and 0.2% penicillin/streptomycin (Gibco, 15140122). Cultures were maintained at 37 °C in a humidified atmosphere of 5% CO₂, with half-medium changes every 3-4 days. All experiments were performed on days *in vitro* 18 (DIV18).

### Reagents

Active human α-synuclein pre-formed fibrils (PFFs), including Type I, ATTO594-conjugated Type I, and Type II (StressMarq Biosciences; SPR-322, SPR-322-A594, and SPR-317, respectively), were diluted in sterile PBS and sonicated for 10 min prior to use. Neurons were treated with PFFs at a final concentration of 5 μg/mL beginning at DIV4 for 14 days, with experiments performed at DIV18. A single treatment of unlabeled Type I PFFs was used unless otherwise indicated (**Fig. S1A**). In control experiments, neurons received an equivalent volume of PBS. TAT-HA-C1aB (CCAD) and the corresponding scrambled control peptide TAT-HA-C1aB-Scr (SCRBL) (peptide sequences provided in Supplementary table 1) were dissolved in sterile molecular biology-grade water and applied at a final concentration of 1 μM for 24 h. Roscovitine (Sigma, R7772) was dissolved in DMSO and applied at a final concentration of 10 μM for 24 h, control experiments included an equivalent volume of DMSO. Tetrodotoxin (TTX; Alomone, T-550), D-AP5 (Alomone, D-145), and CNQX (Alomone, C-141) were dissolved in sterile molecular biology-grade water and applied for 2 h at final concentrations of 1 μM, 10 μM, and 50 μM, respectively. Nimodipine (Alomone, N-150) was dissolved in DMSO and applied at a final concentration of 10 μM for 90 s.

### Western blot

Neurons were scraped in RIPA lysis buffer (Thermo Scientific, 89900) containing Complete/Mini/EDTA-free protease inhibitor cocktail (Roche, 11836170001), sodium fluoride (1 mM; Sigma-Aldrich, 67414), SDS (0.05%), and microcystin (4 μg/mL; Sigma Millipore, 475821), and centrifuged at 13,200 rpm for 20 min at 4 °C. Protein concentration was quantified using the Pierce BCA protein assay kit (Thermo Scientific, 23225) according to the manufacturer’s instructions. 15 μg of protein sample was treated with 2x Laemmli sample buffer (Bio-Rad, 1610737) supplemented with 5% β-mercaptoethanol, and separated on a 5–12% polyacrylamide gel using Tris–glycine running buffer and electrophoresed for approximately 120 min at 125 V. Proteins were transferred onto methanol-activated PVDF membranes (Invitrogen, LC2005) using a Bio-Rad transfer system (1645052) overnight at 25 V. Membranes were blocked in Tris-buffered saline (TBS) supplemented with 0.1% Tween-20 (TBS-T) and 5% non-fat dry milk (hereafter referred to as blocking buffer) for 1 h at RT. Membranes were incubated O/N at 4 °C in blocking buffer with the following primary antibodies: Ca_V_1.2 at 1:250 (Alomone Labs, acc-003), K_V_2.1 at 4 μg/mL (NeuroMab, K89/34), or GAPDH at 1:1000 (Proteintech, 10494-I-AP). After washing with blocking buffer (3 × 10 min), membranes were incubated in blocking buffer with fluorescent secondary antibodies goat anti-rabbit IgG 800CW at 1:10,000 (LI-COR, 926-32211) and goat anti-mouse IgG 680RD at 1:5,000 (LI-COR, 926-68070). Membranes were washed 3 × 10 min with TBS-T. Signals were detected using a Bio-Rad ChemiDoc imaging system and quantified using ImageJ. Protein abundance was normalized to GAPDH.

### Immunocytochemistry

Unless otherwise specified (see c-Fos expression assay), fixation and immunolabeling procedures were consistent throughout the study. Cultured neurons were fixed in 3% paraformaldehyde (PFA) and 0.1% glutaraldehyde (GA) in PBS for 15 min at room temperature (RT). Cells were washed 5 × 5 min with PBS, then blocked and permeabilized in PBS containing 20% Sea Block (Thermo Scientific, 37527) and 0.25% Triton X-100 (Sigma, T8787) for 1 h at RT. Primary antibodies were diluted in the same blocking/permeabilization solution and incubated overnight at 4 °C (antibody details, including dilutions, catalog numbers, and clone identifiers, are provided in Supplementary table 1). Following 5 × 5 min PBS washes, cells were incubated with the appropriate mouse IgG subclass-specific and/or species-specific fluorescent secondary antibodies diluted in blocking/permeabilization solution for 1 h at RT, protected from light. After 5 × 5 min PBS washes, coverslips were mounted on microscope slides using DAPI Fluoromount-G (SouthernBiotech, 0100-20) or retained in PBS for subsequent imaging.

### Confocal microscopy

Images were acquired on one of two systems: (1) a Zeiss LSM880 confocal laser scanning microscope equipped with a Plan-Apochromat 63×/1.40 Oil DIC M27 objective and a super-resolution Airyscan detection unit, or (2) an Olympus FluoView FV4000 confocal laser scanning microscope equipped with a Plan-Apochromat 60×/1.50 Oil HR objective, a UPLXAPO 20×/0.80 objective, and a silicon-based SilVIR detector. The specific system used for each experiment is indicated in the corresponding figure legends. On the Zeiss LSM880, fluorescence signals from DAPI, Alexa Fluor 488, Alexa Fluor 568, and Alexa Fluor 647 were excited using 405-, 488-, 594-, and 633-nm laser lines, respectively. On the Olympus FV4000, the corresponding excitation wavelengths were 405, 488, 561, and 640 nm. Image acquisition was performed using Zeiss ZEN software v2.3 or Olympus cellSens FV software, respectively. Images were acquired as either single optical sections at the internal or PM focal plane, or as Z-stacks collected at axial step sizes of 0.5 μm (Zeiss LSM880) or 0.09 μm (Olympus FV4000). Z-stacks were compressed into maximum intensity projections using ImageJ (NIH). Specific acquisition parameters for each experiment are detailed in the figure legends.

### Super resolution TIRF microscopy

Coverslips were mounted onto glass depression slides (neoLab, Heidelberg, Germany) filled with GLOX-MEA oxygen scavenging buffer (50 mM Tris pH 8.0, 10 mM NaCl, 10% w/v glucose, 0.56 mg/mL glucose oxidase, 34 μg/mL catalase, and 10 mM cysteamine). Coverslips were sealed with Twinsil dental glue (Picodent, Wipperfürth, Germany) and aluminum tape (Thorlabs, T205-1.0-AT205).

Single-molecule localization microscopy was performed on a Leica Infinity TIRF inverted microscope equipped with a 163×/1.49 NA TIRF oil immersion objective and a Hamamatsu ORCA-Flash 4.0 camera. Alexa Fluor 647 was excited using the 638 nm laser line. For each field of view, 50,000 frames were acquired with an exposure time of 10 ms. Single-molecule localization maps were reconstructed with a pixel size of 20 nm. Image acquisition and localization processing were performed using Leica LAS X software.

### Proximity Ligation Assay

The Duolink *In Situ* Proximity Ligation Assay (PLA) kit was used to quantify Ca_V_-K_V_2.1 and K_V_2.1-K_V_2.1 proximity. Neurons were fixed as described in the Immunocytochemistry section, followed by incubation with 100 mM glycine for 15 min at RT. Cells were washed 2 × 5 min in PBS, then blocked and permeabilized in PBS containing 20% Sea Block and 0.25% Triton X-100 for 1 h at RT. Primary antibody pairs were applied in the same solution overnight at 4 °C. The following combinations were used: K_V_2.1 (NeuroMab, K89/34) with Ca_V_1.2 (Alomone, ACC-003); K_V_2.1 (NeuroMab, K89/34) with Ca_V_2.1 (Alomone, ACC-001); and K_V_2.1 (NeuroMab, K89/34) with K_V_2.1 (NeuroMab, Drk1). For negative controls, Ca_V_1.2 antibody (Alomone, ACC-003) was applied alone. Additional antibody details are provided in Supplementary table 1.

Following 2 × 5 min washes with Duolink Wash Buffer A (DUO82049), oligonucleotide-conjugated secondary antibodies (anti-mouse MINUS, DUO92004; anti-rabbit PLUS, DUO92002) were applied at 1:5 dilution in Duolink Antibody Diluent for 1 h at 37 °C. PLA detection was performed using the Duolink *In Situ* Detection Reagents Orange kit (DUO92007) according to the manufacturer’s protocol. Briefly, after 2 × 5 min washes with Wash Buffer A, ligase was applied at 1:40 dilution in Ligation Buffer for 30 min at 37 °C. Following 2 × 5 min washes with Wash Buffer A, polymerase was applied at 1:80 dilution in Amplification Buffer for 100 min at 37 °C, protected from light. Final washes consisted of 2 × 10 min in 1× Wash Buffer B (DUO82049) followed by 1 min in 0.01× Wash Buffer B. Coverslips were mounted with DAPI Fluoromount-G.

PLA images were acquired using either the Zeiss LSM880 or Olympus FV4000 systems described above (see figure legends for details). DAPI and PLA signals were visualized using 405- and 488-nm laser lines, respectively. Z-stacks were collected at axial step sizes of 0.5 μm (Zeiss LSM880) or 0.09 μm (Olympus FV4000) and compressed into maximum intensity projections using ImageJ.

### Live cell cytosolic Ca^2+^ imaging

Neurons were incubated in regular Ringer’s solution (10 mM HEPES pH 7.4, 160 mM NaCl, 2.5 mM KCl, 2 mM CaCl₂, 1 mM MgCl₂, and 8 mM D-glucose) supplemented with 2.5 μM Fluo-4 AM (Invitrogen, F14201) and 0.1% Pluronic F-127 (Thermo Fisher, P3000MP) for 20 min at RT. Cells were then transferred to Fluo-4 AM– and Pluronic F-127–free Ringer’s solution for 10 min at RT to allow de-esterification of the dye.

For KCl depolarization experiments, Fluo-4-loaded neurons were placed in a perfusion chamber with continuous flow of regular Ringer’s solution and imaged on an inverted Andor W1 spinning-disk confocal microscope equipped with an Olympus UPlanFL N 40× oil objective and a Photometrics Prime 95B camera. Fluo-4 fluorescence was excited at 488 nm and acquired at 10 s intervals using Micro-Manager software (v1.4.21). Depolarization was induced by switching to high-KCl Ringer’s solution (10 mM HEPES pH 7.4, 125 mM NaCl, 40 mM KCl, 2 mM CaCl₂, 1 mM MgCl₂, and 8 mM D-glucose) for 50 s.

For electrical field stimulation experiments, Fluo-4-loaded neurons were placed in a field stimulation chamber (Warner Instruments, RC-47FSLP) with continuous flow of regular Ringer’s solution. Stimulation was delivered at 40 V and 1 Hz for 5 s using a MyoPacer field stimulator (IonOptix). Data were acquired at 10 ms intervals.

Intracellular Fluo-4 fluorescence changes were measured within regions of interest (ROIs) placed over the soma, nucleus, or whole-cell soma, as specified in the corresponding figure legends.

### c-Fos expression assay

Neurons were incubated in 1 mL regular Krebs-Ringer buffer (20 mM HEPES pH 7.4, 146 mM NaCl, 4.7 mM KCl, 2.5 mM CaCl₂, 0.15 mM NaH₂PO₄, 1.6 mM NaHCO₃, 0.6 mM MgSO₄, and 8 mM glucose) supplemented with 1 μM TTX, 10 μM D-AP5, and 50 μM CNQX for 2 h at 37 °C to silence synaptic and intrinsic activity. Neurons were then stimulated by addition of 1 mL high-KCl Krebs-Ringer buffer (20 mM HEPES pH 7.4, 62.2 mM NaCl, 90 mM KCl, 2.5 mM CaCl₂, 0.15 mM NaH₂PO₄, 1.6 mM NaHCO₃, 0.6 mM MgSO₄, and 8 mM glucose) supplemented with 1 μM TTX, 10 μM D-AP5, and 50 μM CNQX, yielding a final KCl concentration of 47 mM while maintaining osmolarity at approximately 343 mOsm. For Ca_V_ channel inhibition experiments, the high-KCl buffer additionally contained 10 μM nimodipine. Stimulation was performed for 90 s at 37 °C, after which the buffer was replaced with conditioned culture medium for 3 h at 37 °C to allow c-Fos expression.

Neurons were fixed in PBS containing 4% PFA and 4% (w/v) sucrose for 15 min at 4 °C, washed 3 × 5 min in PBS, and blocked and permeabilized in BLOTTO (10 mM Tris pH 7.4, 150 mM NaCl, 4% w/v non-fat dry milk, and 0.1% Triton X-100) for 1 h at RT. Primary antibodies were diluted in BLOTTO and incubated for 1 h at RT (details provided in Supplementary table 1). Following 3 × 5 min washes with BLOTTO, cells were incubated with the appropriate mouse IgG subclass-specific and/or species-specific fluorescent secondary antibodies diluted in BLOTTO for 1 h at RT, protected from light. After 3 × 5 min PBS washes, coverslips were mounted using DAPI Fluoromount-G.

Images were acquired on the Olympus FV4000 equipped with the UPLXAPO 20×/0.80 objective and SilVIR detector. Fluorescence signals from DAPI, Alexa Fluor 488 (MAP2), and Alexa Fluor 647 (c-Fos) were excited using 405-, 488-, and 640-nm laser lines, respectively. Images were analyzed using ImageJ.

### Image processing, data analysis and figure preparation

All image processing and quantitative analysis were performed using ImageJ. For measurements of cluster size, cluster density, integrated density, and mean gray value (MGV), analysis parameters including background subtraction, thresholding, and particle detection settings were held constant across all conditions within each experiment. Cluster counts were normalized to the respective somatic area. Spatial overlap between two proteins was quantified by multiplying binary masks of each channel to identify colocalized regions. Nearest-neighbor distance analysis was performed using Imaris software (Oxford Instruments).

For live-cell Ca^2+^ imaging experiments, Fluo-4 fluorescence intensity within each ROI was normalized to the minimum intensity value recorded before stimulation. Area-under-the-curve measurements for KCl stimulation experiments were calculated using GraphPad Prism (GraphPad Software). Ca^2+^ peak amplitudes during electrical field stimulation were analyzed using Clampfit software (Molecular Devices).

All statistical analyses were performed using GraphPad Prism. Normality was assessed using the D’Agostino-Pearson omnibus test. Comparisons between two independent groups were made using unpaired two-tailed Student’s t-test (parametric) or Mann-Whitney U-test (non-parametric). For experiments involving four groups with two categorical independent variables, two-way ANOVA was performed followed by appropriate post hoc tests for pairwise comparisons. Statistical significance was defined as P < 0.05 and denoted as follows: *P ≤ 0.05, **P ≤ 0.01, ***P ≤ 0.001, ****P ≤ 0.0001. All data are presented as mean ± SEM. Each experiment was performed using a minimum of two independent cultures; sample sizes are provided in the corresponding figure legends. Schematic illustrations were created using BioRender (BioRender, Inc.).

## Data availability

The data that support the findings of this study are available from the corresponding author upon reasonable request.

## Code availability

Custom analysis codes are available from the corresponding author upon reasonable request.

## Acknowledgements

we are grateful to all members for the Dickson and Dixon laboratories for their helpful input and advice. We thank Joshua Tulman for help generating the graphical abstract. We thank all laboratories for sharing key resources. This work was supported by NIH grants R35 GM149211 and RF1 NS131379 (E.J.D), and NIH grants R01 AG063796 and R01 HL159304 (R.E.D).

## Notes

### Competing Interest Statement

The authors have declared no competing interest.

