## Supplementary Information for "Nanoscale CaV channel reorganization links α-synuclein pathology to calcium-dependent transcriptional dysregulation"

Information within this file:

**Supplementary Fig. 1** Neuronal PFF uptake, cell-type identification, assessment of CaV1.2 and KV2.1 expression, and effects of Type II PFF on CaV1.2 clustering

**Supplementary Fig. 2** CCAD peptide does not alter K_V_2.1 clustering and PLA specificity controls

**Supplementary Fig. 3** PFF treatment does not alter CDK5 expression or subcellular distribution, and PLA specificity controls for roscovitine experiments

**Supplementary Fig. 4** Resting Ca²⁺ levels are comparable between CTL and PFF-treated neurons, but PFF-treated neurons exhibit elevated nuclear Ca^2+^ upon stimulation

**Supplementary table 1:** Resources table

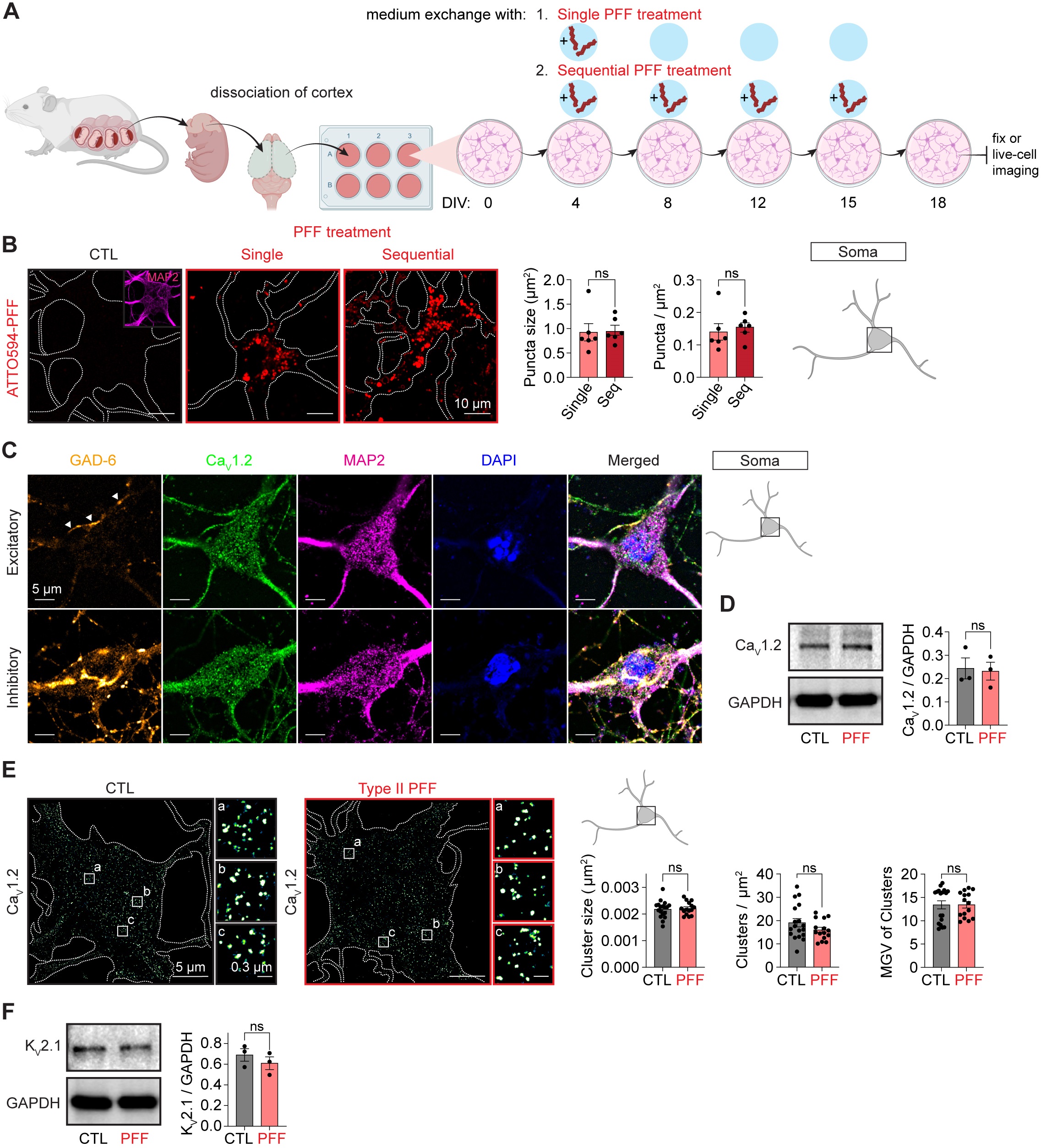

**Supplementary Fig. 1 Neuronal PFF uptake, cell-type identification, assessment of Ca_V_1.2 and K_V_2.1 expression, and effects of Type II PFF on Ca_V_1.2 clustering. A** Schematic illustration of culturing cortical neurons and design of PFF treatment. **B** Left: representative single-plane Airyscan confocal images of neuronal cross-sections showing ATTO594-PFF fluorescence in PFF-treated (red) neurons. CTL (black) neurons were counterstained for MAP2 (pink) to delineate cell boundaries (inset). In the single-treatment paradigm, ATTO594-PFFs were applied only at DIV4, and the medium was changed every 3–4 days without additional PFF. In the sequential-treatment paradigm, ATTO594-PFFs were added at DIV4 and replenished with each medium change throughout the 14-day treatment period. Right: quantification of ATTO594-PFF puncta size and puncta density in single (light red) and sequential (Seq, dark red) PFF-treated neurons. n = 6 somata per condition; one isolation. **C** Representative single-plane Airyscan confocal images of the PM showing immunolabeling for GAD-67 (inhibitory neuronal marker), Ca_V_1.2, MAP2 (neuronal marker), and DAPI (nucleus) in CTL excitatory and inhibitory neurons. White arrows indicate inhibitory neuron projections surrounding the excitatory neuron soma. **D** Representative western blots from control and PFF-treated cultured neuron lysates probed for Ca_V_1.2 and GAPDH. Three isolations. **E** Left: representative super-resolution TIRF localization maps showing Ca_V_1.2 immunolabeling in CTL (black) and Type II PFF-treated (red) neurons. Right: quantification of PM Ca_V_1.2 cluster size, cluster density, and mean gray value (MGV) in the somatic region. n = 18 (CTL) and n = 15 (Type II PFF) neurons; two independent isolations. **F** Same as (**D**) only for K_V_2.1. Three isolations. Error bars represent SEM. Statistical significance in panels (**B**) and (**D-F**) was determined using two-tailed Mann-Whitney or unpaired two-tailed t-tests. ns, not significant. CTL, control; Type II PFF, α-synuclein pre-formed fibril Type II treatment; GAD-6, monoclonal antibody for glutamic acid decarboxylase 67 (inhibitory neuronal marker); MAP2, microtubule-associated protein 2 (neuronal marker).

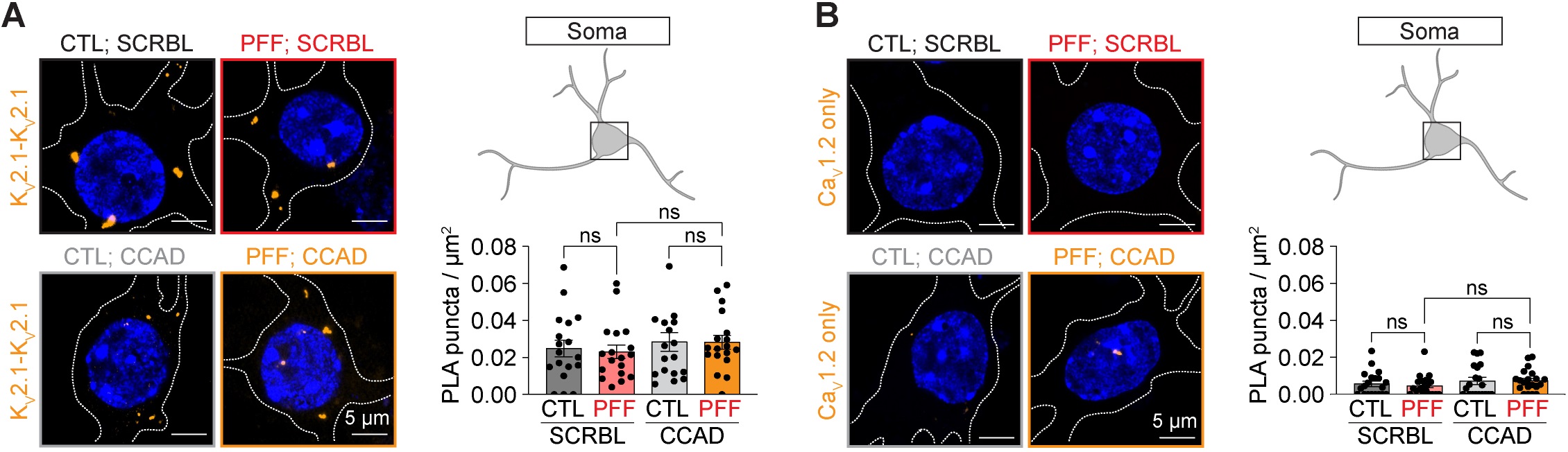

**Supplementary Fig. 2 CCAD peptide does not alter K_V_2.1 clustering and PLA specificity controls.** **A** Left: representative Airyscan confocal PLA images showing K_V_2.1-K_V_2.1 proximity in CTL;SCRBL (black), PFF;SCRBL (red), CTL;CCAD (gray), and PFF;CCAD (yellow) neurons. Images are maximum intensity projections from Z-stacks spanning whole cells. Right: quantification of PLA puncta density. n = 18 neurons per condition; two independent isolations. **B** PLA specificity control using only a single Ca_V_1.2 primary antibody. Left: representative Airyscan confocal PLA images. Color coding as in (**A**). Right: quantification of PLA puncta density. n = 20 neurons per condition; two independent isolations. Error bars represent SEM. Statistical significance in panels (**A**–**B**) was determined using two-way ANOVA with appropriate post hoc tests. ns, not significant. CTL, control; PFF, α-synuclein pre-formed fibril treatment; CCAD, calcium channel association domain peptide; SCRBL, scrambled control peptide.

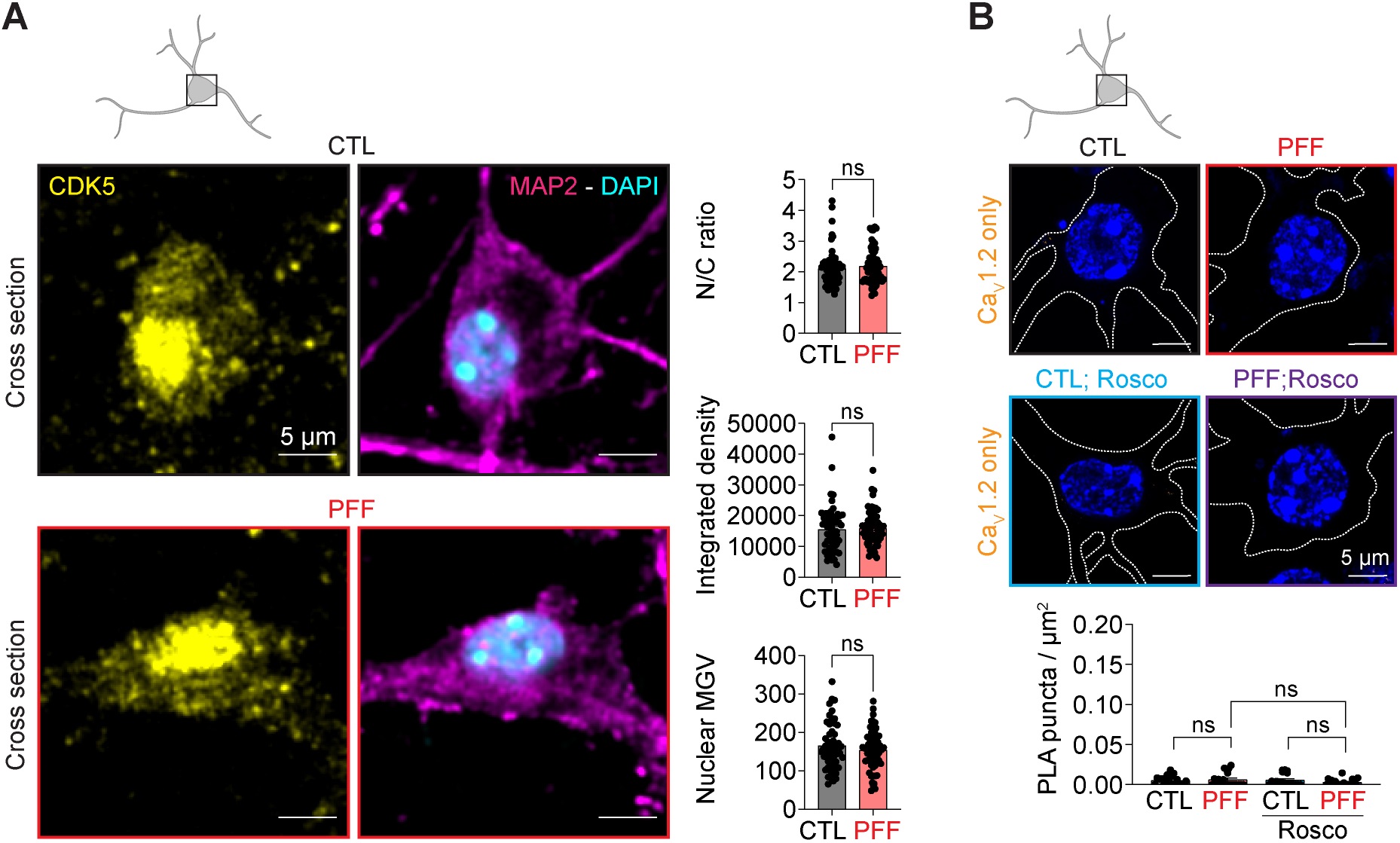

**Supplementary Fig. 3 PFF treatment does not alter CDK5 expression or subcellular distribution, and PLA specificity controls for roscovitine experiments. A** Left: representative single-plane FV4000 confocal images showing CDK5 immunolabeling in cross-sections of CTL (black) and PFF-treated (red) neurons. DAPI and MAP2 were used to identify nuclei and cell boundaries, respectively. Right: quantification of CDK5 nuclear-to-cytoplasmic (N/C) ratio, integrated density, and nuclear mean gray value (MGV) in CTL (black) and PFF-treated (red) neurons. n = 59 (CTL) and n = 64 (PFF) neurons; two independent isolations. **B** PLA specificity control using only a single Ca_V_1.2 primary antibody in the roscovitine experimental paradigm. Left: representative FV4000 confocal PLA images in CTL (black), PFF (red), CTL;Rosco (blue), and PFF;Rosco (purple) neurons. Images are maximum intensity projections from Z-stacks spanning whole cells. Right: quantification of PLA puncta density. n = 20 neurons per condition; two independent isolations. Error bars represent SEM. Statistical significance in panel (**A**) was determined using two-tailed Mann-Whitney or unpaired two-tailed t-tests; panel (**B**) was analyzed using two-way ANOVA with appropriate post hoc tests. ns, not significant. CTL, control; PFF, α-synuclein pre-formed fibril treatment; CDK5, cyclin-dependent kinase 5; MAP2, microtubule-associated protein 2 (neuronal marker); Rosco, roscovitine.

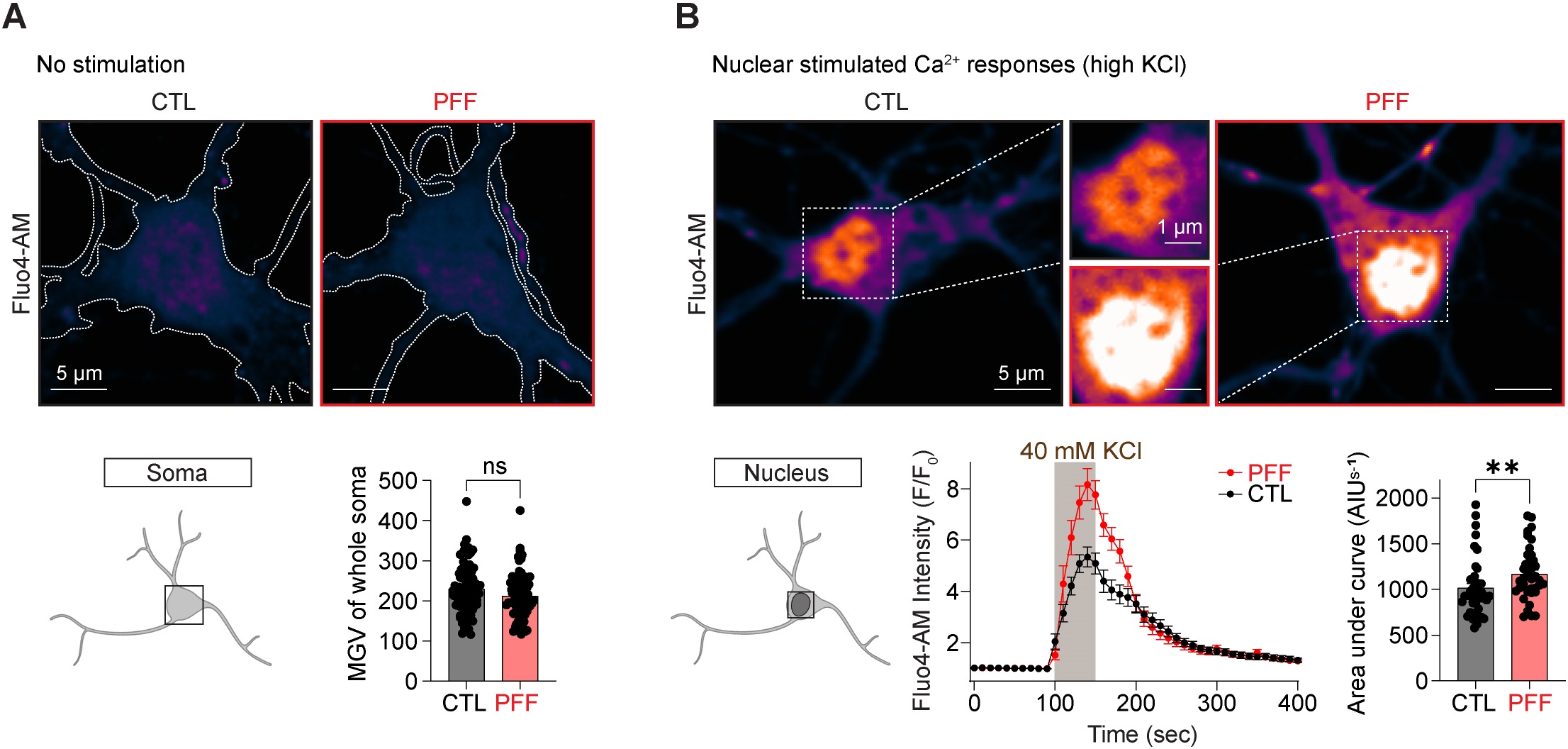

**Supplementary Fig. 4 Resting Ca²⁺ levels are comparable between CTL and PFF-treated neurons, but PFF-treated neurons exhibit elevated nuclear Ca^2+^ upon stimulation.** **A** Top: representative live-cell spinning-disk confocal single-plane images of Fluo-4 AM-loaded CTL (black) and PFF-treated (red) neurons in the absence of stimulation. Bottom: quantification of whole-soma mean gray value (MGV) in CTL (black) and PFF-treated (red) neurons. MGV was calculated as the average of 20 s continuous recordings acquired at 50 ms per frame. n = 88 (CTL) and n = 75 (PFF) neurons; two independent isolations. **B** Top: representative live-cell spinning-disk confocal single-plane images of Fluo-4 AM-loaded CTL (black) and PFF-treated (red) neurons upon stimulation with 40 mM KCl. Bottom, middle: averaged and normalized Fluo-4 AM fluorescence time series showing nuclear Ca²⁺ responses to 40 mM KCl stimulation (light gray shading) in CTL (black) and PFF-treated (red) neurons. Bottom, right: quantification of area under curve from normalized nuclear Fluo-4 AM intensity traces. n = 42 (CTL) and n = 39 (PFF) nuclei; two independent isolations. Error bars represent SEM. Statistical significance was determined using two-tailed Mann-Whitney tests. ns, not significant; ****P ≤ 0.0001. CTL, control; PFF, α-synuclein pre-formed fibril treatment.

**Supplementary table 1: Resources table**

| **REAGENT or RESOURCE** | **SOURCE** | **IDENTIFIER** |
| --- | --- | --- |
| **Antibodies** |  |  |
| Rabbit anti-CaV1.2 (1:300 for IF and 1:250 for WB) | Alomone Labs | Cat # ACC-003 RRID:AB_2039771 |
| Rabbit anti-CaV2.1 (1:200 for IF) | Alomone Labs | Cat # ACC-001 RRID:AB_2039764 |
| Mouse anti-Kv2.1 IgG2a (1:20 for IF) | UC Davis/NIH NeuroMab Facility | Cat # K89/34R-rat IgG2a, RRID:AB_3112073 |
| Mouse anti-Kv2.1 IgG1 (5 µg/mL for IF and 4 µg/mL for WB) | UC Davis/NIH NeuroMab Facility | Cat # K89/34 RRID:AB_2877280 |
| Rabbit anti-Kv2.1 (1:100) | UC Davis/NIH NeuroMab Facility | Cat# DRK1, RRID:AB_2891234 |
| Mouse anti-p(603)Kv2.1 IgG1 (1:5 for IF) | UC Davis/NIH NeuroMab Facility | Cat # L61/14 RRID:AB_2315769 |
| Mouse anti-RyR IgG1 (1:100 for IF) | Abcam | Cat # Ab2868 RRID:AB_2183051 |
| Chicken anti-MAP2 IgY (1:1000 for IF) | Abcam | Cat # ab5392 RRID:AB_2138153 |
| Mouse anti-GAD65/67 IgG2a (1:1.5 for IF) | UC Davis/NIH NeuroMab Facility | Cat# L127/12, RRID:AB_2756510 |
| Mouse anti-C-fos IgG1 (1:5 for IF) | UC Davis/NIH NeuroMab Facility | Cat# N486/76, RRID:AB_2833039 |
| Mouse anti-CDK5 IgG1 (1:50 for IF) | Santa Cruz Biotechnology | Cat# sc-6247, RRID:AB_627241 |
| Rabbit anti-GAPDH (1:1000 for WB) | Proteintech | Cat# 10494-1-AP, RRID:AB_2263076 |
| Goat anti-chicken IgY (AF488) (1:1000 for IF) | Invitrogen | Cat # A32931 RRID:AB_2762843 |
| Goat anti-mouse IgG1 (AF568) (1:1000 for IF) | Invitrogen | Cat # A21124 RRID:AB_2535766 |
| Goat anti-mouse IgG1 (AF647) (1:1000 for IF) | Invitrogen | Cat # A21240 RRID:AB_2535809 |
| Goat anti-mouse IgG2a (AF568) (1:1000 for IF) | Invitrogen | Cat # A21134 RRID:AB_1500825 |
| Goat anti-rabbit IgG (AF647) (1:1000 for IF) | Invitrogen | Cat # A21245 RRID:AB_2535813 |
| goat anti-rabbit IgG 800CW (1:10,000 for WB) | LI-COR | Cat # 926-32211 RRID:AB_621843 |
| goat anti-mouse IgG 680RD (1:5,000 for WB) | LI-COR | Cat # 926-68070 RRID:AB_10956588 |
| **Recombinant proteins, Peptides, and Chemicals** |  |  |
| Active human α-synuclein pre-formed fibrils (PFFs) Type I (5 μg/mL, 14 days) | StressMarq Biosciences | SPR-322 |
| Active human α-synuclein pre-formed fibrils (PFFs) ATTO594-conjugated Type I (5 μg/mL, 14 days) | StressMarq Biosciences | SPR-322-A594 |
| Active human α-synuclein pre-formed fibrils (PFFs) Type II (5 μg/mL, 14 days) | StressMarq Biosciences | SPR-317 |
| TAT-HA-C1aB (CCAD, 1 µM, 24 h) Seq: GRKKRRQRRRYPYDVPDYAHLSPNKWKW | Genscript | N/A |
| TAT-HA-C1aB-Scr (Scr, 1 µM, 24 h) Seq: GRKKRRQRRRYPYDVPDYANLKWSHPKW | Genscript | N/A |
| Roscovitine (10 µM, 24 h) | Sigma-Aldrich | Cat # R7772 |
| Tetrodotoxin (1 µM, 2 h) | Alomone | Cat # T-550 |
| CNQX (50 µM, 2 h) | Alomone | CaT # C-141 |
| AP5 (10 µM, 2 h) | Alomone | Cat # D-145 |
| Nimodipine (10 µM, 90 sec) | Alomone | Cat # N-150 |
| Fluo-4-AM (2.5 µM, 20 min) | Life Technologies | Cat # F14201 |
| Pluronic F-127 | Thermo Fisher Scientific | Cat # P3000MP |
| Neurobasal medium | Gibco | Cat # 21103-049 |
| B27 | Gibco | Cat # 17504-044 |
| Glutamax | Gibco | Cat # 35050-061 |
| Penicillin/streptomycin | Gibco | Cat # 15140122 |
| Poly-D-lysine | Sigma | Cat # P6407 |
| SEA BLOCK Blocking Buffer | Thermo Scientific | Cat # 37527 |
| Paraformaldehyde | Electron Microscopy Sciences | Cat # 15710 |
| Triton X-100 | Sigma-Aldrich | Cat # T8787 |
| DAPI Fluoromount-G | SouthernBiotech | Cat # 0100-20 |
| DAPI | Sigma-Aldrich | Cat # D9542 |
| RIPA buffer | Thermo Scientific | Cat # 89900 |
| Complete/Mini/EDTA-free protease inhibitor cocktail | Roche | Cat # 11836170001 |
| Sodium fluoride (1 mM) | Sigma-Aldrich | Cat # 67414 |
| Microcystin (4 μg/mL) | Sigma-Aldrich | Cat # 475821 |
| 2x Laemmli sample buffer | Bio-Rad | Cat # 1610737 |
| PVDF/Filter Paper Sandwiches, 0.45 μm, 8.3 x 7.3 cm | Invitrolon | Cat # LC2005 |
| **Kits** |  |  |
| Papain Dissociation System | Worthington | Cat # LK003150 |
| Duolink In Situ Wash Buffers, Fluorescence | Sigma-Aldrich | Cat # DUO82049-4L |
| Duolink In Situ PLA probe Anti-Mouse MINUS | Sigma-Aldrich | Cat # DUO92004-100RXN |
| Duolink In Situ PLA Probe Anti-Rabbit PLUS | Sigma-Aldrich | Cat # DUO92002-100RXN |
| Duolink In Situ Detection Reagents Orange kit | Sigma-Aldrich | Cat # DUO92007 |
| Pierce BCA protein assay kit | Thermo Fisher Scientific | Cat # 23225 |
